# Tracking the conformational dynamics of single molecules in live cells; GTPase cycling in Cdc42 gradients and microclusters

**DOI:** 10.64898/2026.11.09.765066

**Authors:** Pengning Xu, Saygin Gulec, Nicholas K. Pinkin, Yu Yan, Lihua He, Nathan D. Smyers, Pothiappan Vairaprakash, Frederico M. Pimenta, Shiqiong Hu, Bei Liu, Timothy C. Elston, Klaus M. Hahn

## Abstract

The Rho family of GTPases play a critical role in many cellular processes by acting as molecular switches relying on conformational changes to transition between active and inactive states. The spatial distribution and kinetics of these conformational changes are complex and vary across the cell. We describe here biosensors based on environment-responsive fluorescent dyes, capable of reporting the conformational changes of individual GTPase molecules as they traffic within living cells (SPARC Cdc42 and SPARC Rac1, “Single Particle Activation Reporter Constructs”). We use SPARC Cdc42 to determine the rates of different steps in GTPase regulation, activation/inactivation and membrane insertion, that together generate localized gradients of Cdc42 activity driving cell motility. The biosensors also revealed heterogeneously distributed clusters of Cdc42, the mechanisms of their activation, and potential roles in polarized motility.

## Results

Many proteins involved in cell signaling act as switches, adopting different conformations that interact with distinct binding partners. Because cell behavior is controlled by the subcellular location and kinetics of switching, a full understanding of cellular ‘decision making’ depends on visualizing and quantifying conformational changes within living cells. Although this has been accomplished with fluorescent biosensors based on a host of different designs, the light output and signal/noise ratio of these biosensors has limited them in most cases to reporting on protein populations rather than single protein molecules (*1, 2*). Seeing the conformational changes of single molecules in live cells would allow measurement of the spatiotemporal dynamics of signaling networks with submicron precision and reveal mechanisms that underlie the self-organization of subcellular structures.

We designed biosensors for Rho family GTPase enzymes (Fig. 1A), molecular switches that interact with downstream effectors only when they bind GTP and adopt an activated conformation. We sought to produce fluorescent GTPase analogs with sufficient light output to track individual molecules, but also with conformation-dependent emission changes that could simultaneously report conformation. Biosensors for the GTPases Cdc42 and Rac1 were made by linking each to an “affinity reagent” (AR), a protein fragment that bound selectively to the activated GTPase, as in previous successful biosensor designs. Here we added a SNAP domain (SNAP = a fragment of the O⁶-alkylguanine-DNA alkyltransferase enzyme that catalyzes self-labeling by appropriately derivatized small molecules) to the linker between the AR and the GTPase, enabling incorporation of a solvent-sensitive fluorescent dye that responded to changes in its environment when the AR bound the activated GTPase (*3*). The dye was directly excited, unlike in FRET and other biosensors, producing a bright signal, and could be introduced simply by incubating membrane-permeable dye in the extracellular medium. GTPase binding regions, Cdc42 binding domain (CBD) and Pak1 binding domain (PBD), from two different downstream GTPase effectors were combined to generate AR optimized for Rac1 or Cdc42 (Fig. S1A-E) (*4–7*).

**Figure 1.**
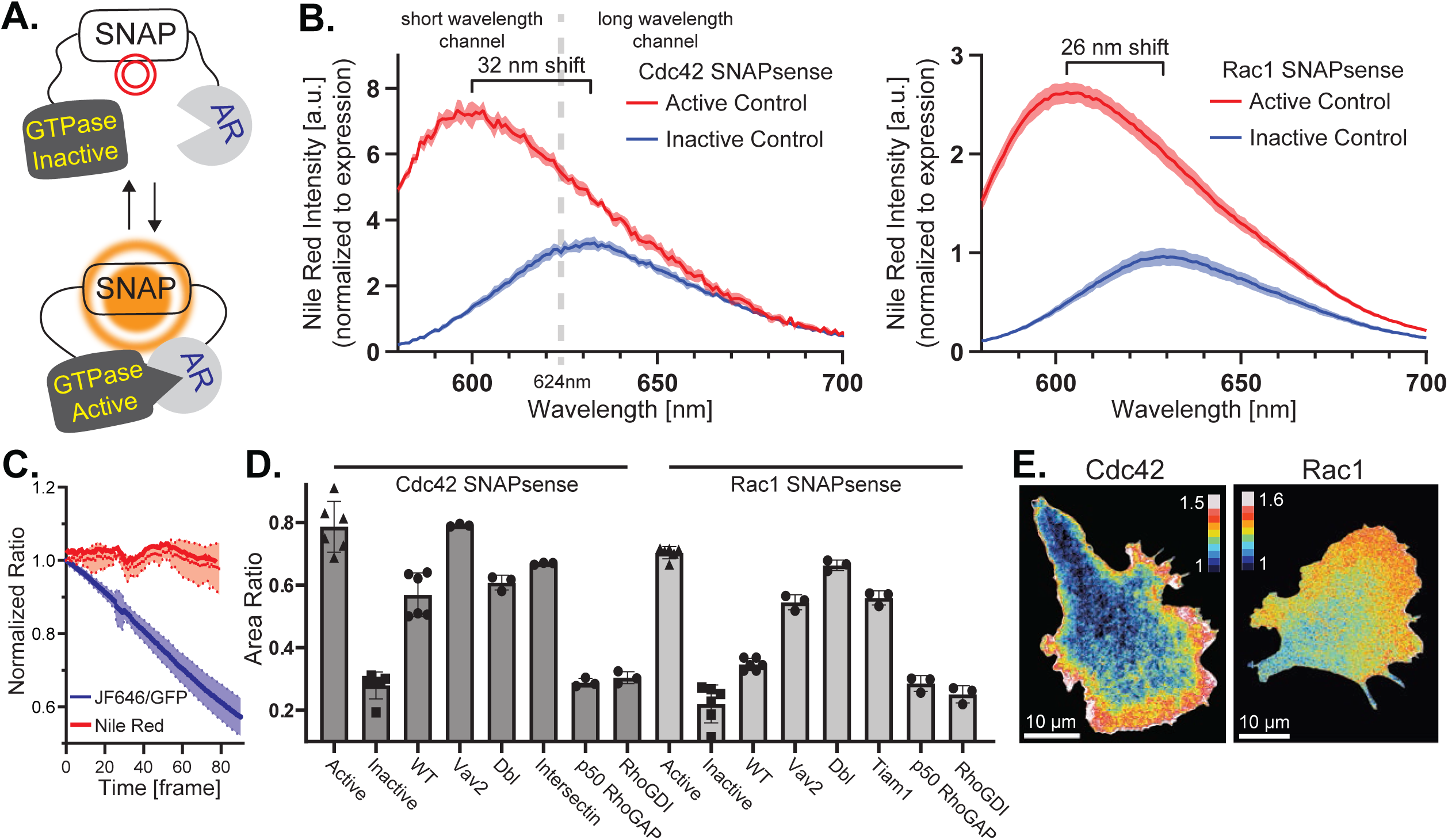
SPARC biosensors. (A) Single Particle Activation Reporter for Cells (SPARC). A peptide “affinity reagent” (AR) is connected to the GTPase via a linker that contains a SNAP domain. Binding of the affinity reagent to the activated conformation of the GTPase changes the environment of the dye, causing a shift in its fluorescence emission. (B) Emission spectra of Cdc42 and Rac1 SPARC biosensors. Curves show the difference between the constitutively active G12V mutant and the inactive biosensor, made by introducing a T17N mutation in the GTPase and two point mutations in the AR. Nile Red intensities are normalized to attached GFP. (C) Ratiometric imaging without bleach correction. Ratio changes during time-lapse imaging; Compare the ratio of two different emission bands from Nile Red (red) to the JF646 dye / sfGFP ratio from a dye-labeled SNAP-sfGFP fusion (blue). (D) SPARC biosensor response to upstream regulators. Emission ratios of biosensors co-expressed with GEFs, GAPs and GDI are shown. GTPase specificity and effects on activity were like wild type GTPases. Active and inactive controls from B are shown for comparison. (E) TIRF imaging of MEF cells expressing SPARC Cdc42 (left) or SPARC Rac1 (right), showing elevated activity at the cell edge.

We made a SNAP-reactive version of Nile Red, a dye with a long history of applications sensing hydrophobic environments in cells (Fig. S1F, G, Supplemental Table 1) (*8–10*). Experimentation with the position of the SNAP-reactive group on Nile Red and the linkers between protein domains led to biosensors for Rac1 and Cdc42 that showed a substantial blueshift and intensity increase upon GTPase activation (Fig. 1B and S1A). We examined the biosensors’ behavior in live cells by comparing constitutively active and inactive mutants, and mutations that abrogate GTPase-AR binding, using fluorescence spectra of cell suspensions and imaging of adherent cells. Incubation of the SNAP-reactive Nile Red in the extracellular medium of mouse embryo fibroblast cells (MEF) expressing biosensor led to good covalent labeling with low background. Fortuitously, the dye underwent a 48-fold increase in fluorescence intensity upon covalent attachment to the biosensor (Fig. S1H, I), and unattached dye could be effectively removed by washing the cells. Attachment of the biosensor to the membrane did not alter the spectra of active or inactive mutants, showing that the dye was not simply responding to activation-induced translocation to the plasma membrane (Fig. S1J).

The spectral shift upon activation (Fig. 1B) was quantified in cells by monitoring the ratio of light output from two relatively broad wavelength ranges, to maximize light collection (574-624 nm, and 624-680 nm). This ratio underwent a 3-fold change for Cdc42 activation and 2.4-fold change for Rac1 (Fig. 1B). One emission band remained more constant while the other changed intensity greatly in response to activation. Single molecules were readily discernable in either band. Because the dye underwent a ratiometric change, corrections for bleaching were not necessary, unlike biosensors using the ratio of light from two different fluorophores that bleach at different rates (e.g. FRET, Fig. 1C). As a reporter of GTPase activation, it was essential that the biosensor respond to upstream regulatory molecules. Both Cdc42 and Rac1 showed normal responses and specificity upon co-expression of guanine exchange factors (activators), GTPase accelerating proteins (deactivators) and guanine dissociation inhibitor (controlling membrane translocation) (Fig. 1D). We also saw that the biosensor was suitable for both ratiometric imaging (Figs. 1E) and fluorescence lifetime imaging (FLIM), with a 0.4 ns shift in fluorescence lifetime upon activation (Figs. S1K-M). The biosensors were named Cdc42 SPARC and Rac1 SPARC (for single particle activation reporter construct).

We began our characterization of single molecule behavior by recording fluorescence from the two emission bands while tracking Cdc42 molecules (Fig. 2A). We used these recordings to construct a 2D histogram of fluorescence intensities from the two bands (Fig. 2A). This revealed two populations of molecules corresponding to the active and inactive states. The >624 nm emission band remained more constant while molecules diffused on the membrane and, therefore, best revealed the arrival, departure and position of molecules on the membrane (Fig. 2A, red curve). Adoption of an active (inactive) conformation led to an increase (or decrease) in intensity for the emission band below 624 nm (Fig. 2A, green curve).

**Figure 2.**
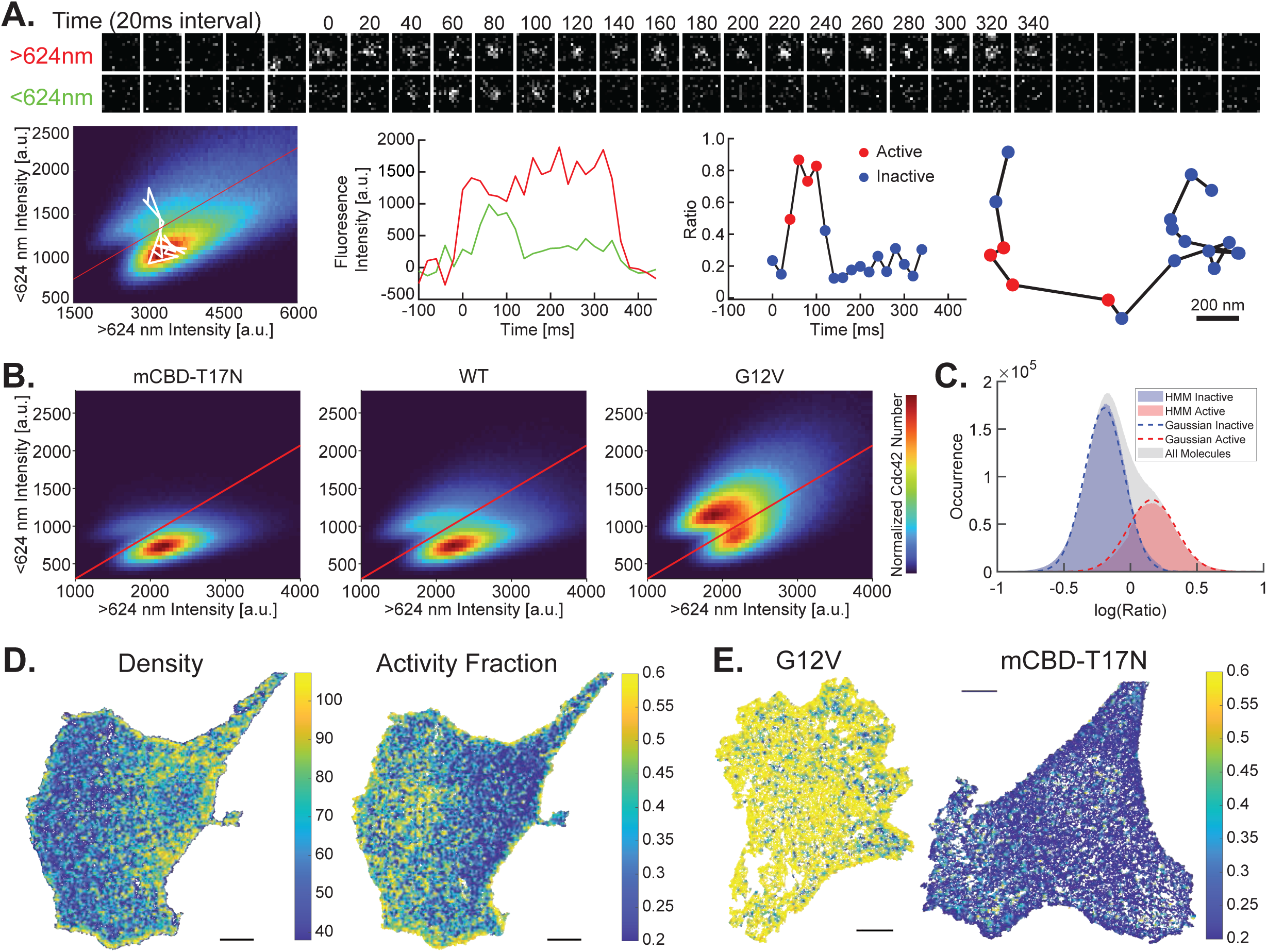
Single-molecule ratio imaging using SPARC Cdc42. (A) Single molecule imaging. Top: Images for a single molecule from the two emission bands. The intensity in the >624 nm band remained relatively constant while the molecule is on the membrane (0-340 ms), whereas intensity from the <624 nm band increased upon activation (40-120 ms). Bottom left to right: 2D histogram of fluorescence measurements from the two emission bands for all single molecule tracks from the same cell. The white curve was generated from the time series shown in the top panel; Fluorescence intensity time series from the two emission bands: >624 nm (red) and <624 nm (green); A time series for the ratio of the two channels for the time series shown in the left panel; The location track for the shown molecule. (B) 2D histograms of intensities from the two emission bands. HeLa cells were transfected with inactivated biosensor mCBD-T17N (N=6), WT biosensor (N=7) and constitutively active G12V (N=6). (C) Histogram of ratios from the two emission bands for WT cells along with distributions found from the HMM for the active and inactive states. (D) Density and activity maps from single molecule tracks, for MEF cells expressing SPARC Cdc42. Each point marks the centroid of a single track. In the density map, color indicates the number of molecules within 500nm. In the activity map, it shows the fraction of active molecules within 500 nm. Scale bar: 10 μm. (E) Single-particle activity maps of MEF cell expressing active (left) and inactive (right) mutants of SPARC Cdc42. Color indicates the fraction of active molecules within 500 nm. Scale bar: 10 μm.

While the 2D histograms and visual inspection of fluorescence intensity along single molecule tracks revealed distinct conformational states, the individual tracks showed significant fluctuations in each of the two emission bands, and the location of the two conformational states within the histograms varied from cell to cell (Fig. S2A). Therefore, we developed an analysis pipeline to track individual molecules, perform local background correction and appropriately scale data sets from different cells to minimize the effects of background fluctuations and cell-to-cell variability (Fig. S2A, B). Following this data processing, the 2D histogram constructed by combining data from all cells produced two clear conformational states and revealed that membrane-bound Cdc42 is primarily in the inactive conformation. This observation, made possible by single molecule imaging, sets limits on the role and timing of activation/deactivation relative to membrane insertion, and will be explored further below by determining the rates of these individual steps. Histograms constructed from inactivated (mCBD-T17N) and constitutively active (G12V) controls showed repressed and elevated molecular activity, respectively (Fig. 2B and S2C).

Control experiments showed that overexpression of the WT biosensor led to increased Cdc42 activity, perhaps because native regulatory pathways were overwhelmed (Fig. S2D) (*11*). For biological studies, the biosensor could be used at well below endogenous concentrations, an advantage of single particle imaging (Fig. S2E). Controls showed that this abrogated overexpression artefacts (Fig. S2D, E).

We observed that the log ratio of fluorescence intensities from the two emission bands was well approximated as the sum of two Gaussian distributions (Fig. 2C). Therefore, to infer the state of individual Cdc42 molecules as they diffused along the cell membrane, we applied a two-state Gaussian hidden Markov Model (HMM) (Fig 2C and S2F, Methods). HMMs are statistical models used to predict unobservable (hidden) states of a system (e.g., protein conformations) from noisy time series data for measurable quantities that are functionally related to the hidden states (e.g., log ratio measurements). HMMs have been used previously to analyze single-molecule data (*12, 13*). The HMM allowed us to assign the most probable conformation state of each molecule at each time point (Fig 2A, bottom two right panels).

Having assigned an activity state to each time point of all the tracks, we used this data to construct maps of Cdc42 distribution and activity in polarized, moving cells. Like traditional biosensors, our maps showed that activated Cdc42 was concentrated at the leading edge, while the bulk of the Cdc42 was found further towards the rear of the cell. Thus, a small percentage of the total Cdc42 population was activated, and this was concentrated at the protruding cell edge where it is known to control actin polymerization (Figs. 2D, S2G, Methods) (*14–17*). In contrast, the inactivated (mCBD-T17N) and active (G12V) controls showed repressed and elevated activity, broadly distributed throughout the cell (Fig. 2E).

Resolving the conformational dynamics of single molecules provided a powerful tool for understanding how localized GTPase activation was produced. We set out to ask how the activation gradients that drove polarized motility were produced by combined regulation of GTPase membrane translocation and localized activation/deactivation (*18*). We first used stochastic modeling to calculate localized transition rates for Cdc42 activation/inactivation and membrane dissociation, and to determine the fraction of molecules that arrive at the membrane in the active state. These were compared for cells expressing WT biosensors, or inactivated and active controls (Figs. 3A, B, Methods). The bleaching of SPARC Cdc42 was more than 10-fold slower than the Cdc42 dissociation rate and allows enough time for observing most Cdc42 tracks (Fig. S3A, B) and has little effect on the kinetic modeling. Previous studies have demonstrated that Cdc42 activity decreases with distance from the cell edge in cell protrusions (*14, 19*). To relate Cdc42 activity to distance from the cell edge, we determined Cdc42 behaviors in different 1-µm layers extending inwards from the cell perimeter (Fig. 3C) (*20–22*). The fraction of time Cdc42 spent in the active state and the Cdc42 molecular density were quantified within each layer. Consistent with previous studies (*15, 19*), layer 1 showed the highest fraction of active molecules, and activity decreased toward the cell center (Fig. 3D, top panel). Molecular density also was elevated in layers 1 and 2 relative to interior layers (Fig. 3D, bottom panel).

**Figure 3.**
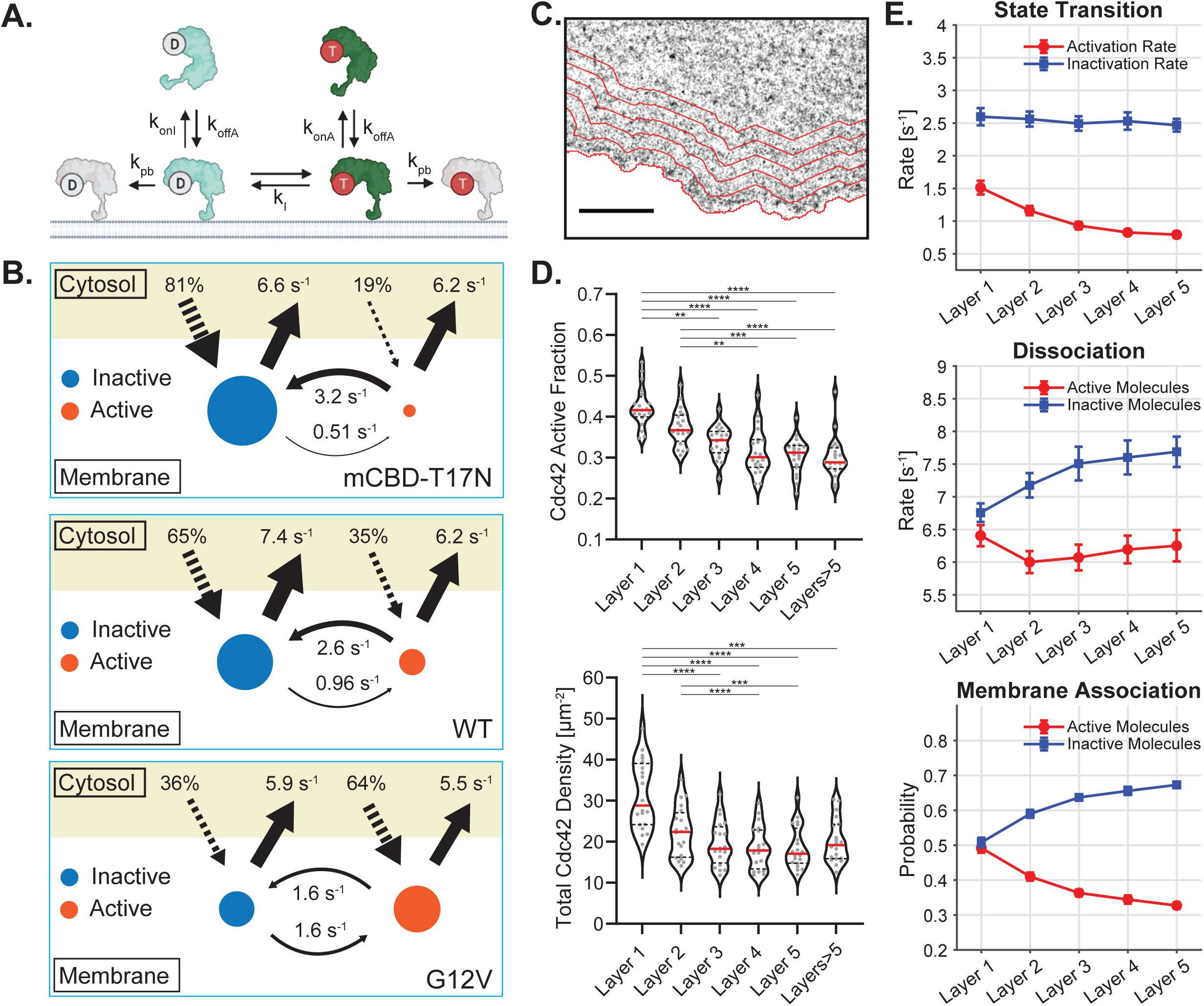
**Chemical kinetic measurements using SPARC**. (A) Schematic diagram for the reactions considered in the stochastic model. (B) Inferred reaction kinetic rates and arrival probabilities from the HMM. (C) Image of a cell edge with layer boundaries denoting distance from the edge shown in red. Scale bar: 5 μm. (D) Cdc42 active fraction (top panel) and total Cdc42 concentration (bottom panel) as a function of distance from the cell edge. (E) Inferred rates for state transitions (top panel), dissociation (middle panel) and association probabilities (bottom panel) as a function of distance from the cell edge.

Because Cdc42 SPARC monitors individual molecules, we were able to determine how the kinetics of Cdc42 membrane dissociation and activation/inactivation varied as a function of distance from the cell edge. Activation rates were more than 50% higher at the edge than at the 5 μm layer, whereas the inactivation rate was similar across all layers (Figure 3E, top panel). The dissociation rate of inactive molecules increased as a function of distance from the edge whereas the dissociation rate of active molecules showed a slight decrease (Fig. 3E, middle panel). The probability of a molecule being active when it arrived at the membrane decreased with distance from the cell edge (Fig. 3E, bottom panel).

Our results suggest that elevated Cdc42 activity near the cell edge is driven by increased activation rates and preferential membrane association of active molecules, rather than by localized suppression of inactivation or changes in membrane dissociation. To further investigate Cdc42 regulation, we developed a reaction-diffusion model for the concentrations of active and inactive Cdc42 molecules based on the schematic diagrams shown in Figs. 3A and 4A (Methods). The inputs for the model were the spatial profiles for the state transition rates, dissociation and association rates, and the association probabilities for arriving at the membrane in the active or inactive state (Fig 4A). Using these inputs the model accurately reproduced the concentration profiles for active and inactive molecules (Fig. 4B, two left panels), from which the total Cdc42 concentration and active fraction could be calculated (Fig. 4B, two right panels). Having demonstrated that the model produced results consistent with our experimental measurements, we used it to predict which of the parameter profiles shown in Fig. 4A had the largest effect on activity gradients. To gauge the relative importance of each input in generating the graded distribution of active Cdc42 molecules, the inputs were each set to their mean values throughout the spatial domain. The association rate had the largest effect on total Cdc42, inactive Cdc42 concentration, and most importantly the concentration of active Cdc42 molecules (Fig. 4C). The probability that active molecules associated with the membrane had the greatest effect on the fraction of total Cdc42 molecules that were active. What appears to be preferential association of active molecules with the membrane could be due to molecules arriving at the membrane in the active state, or membrane translocation and activation may occur concomitantly within the temporal resolution of this study. In conclusion, the gradient of activated Cdc42 molecules, highest at the cell edge, forms primarily because Cdc42 molecules arrive at the membrane in an active state or have a high probability of becoming activated immediately following insertion into the membrane (*23*).

**Figure 4.**
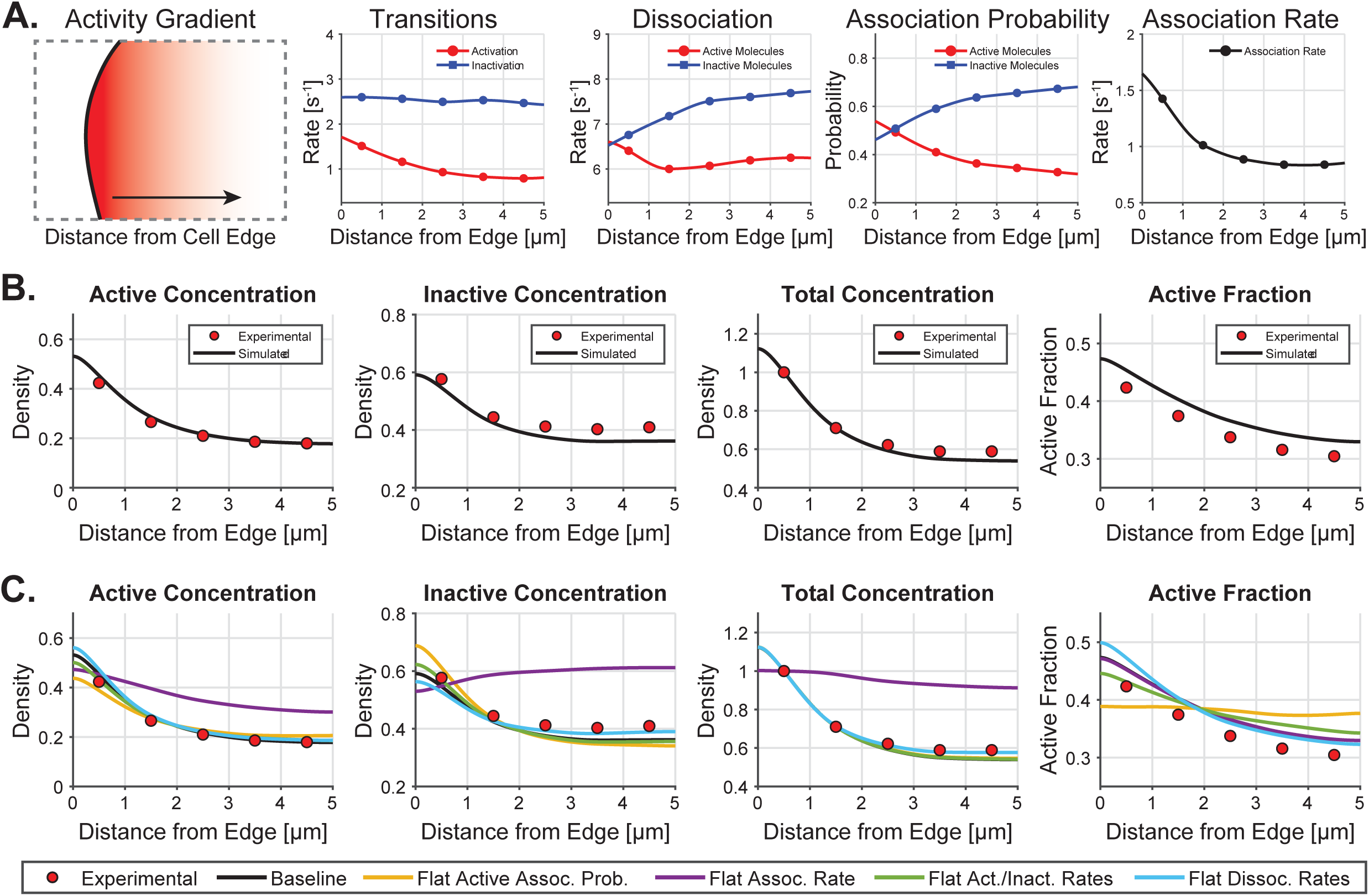
**Reaction-diffusion model for Cdc42 activity**. (A) Schematic diagram for the model (left panel) and the spatial profiles for the reaction rates and association probabilities (right four panels) used as input for the model. (B) Concentrations for active, inactive and total Cdc42 and the active fraction. The black curves are the model results and red circles are the experimental data. (C) Predicted concentration profiles when the spatial gradient for each input rate is removed and replaced with its mean value.

Visual inspection of single-particle tracks suggested the presence of multi-molecule clusters of Cdc42 (Fig. 3C & S4A). Cdc42 nanoclusters previously have been observed in the context of budding and fission yeast and have been suggested to function as a counter measure to dissipative diffusion (*24*). To establish the existence of clusters, we applied a normalized version of the Ripley’s K Function (H(r)) to Cdc42 track centroids. H(r) measures clustering strength as the deviation of the observed pair-wise particle distribution from that of a uniform distribution. This analysis confirmed Cdc42 clustering and estimated a cluster radius of 164 nm (Fig. 5A, Methods), well below the diffraction limit of a light microscope. Photoactivation localization microscopy (PALM) imaging of Cdc42 attached to PAmCherry in fixed MEF cells revealed that both wildtype Cdc42 and constitutively active Cdc42CA were organized in clusters (Fig. S4A) with an average size of 84±13 nm and 79 ± 11 nm, respectively (Fig. S4B). The difference in cluster size between the centroid and PALM measurements might result from dynamic features of clusters not captured by PALM. We used DBSCAN to identify and characterize the clusters from our track data (Methods) (*25*). This procedure distinguishes three types of molecules: those in the core of the clusters, on the border of clusters, and non-clustered molecules. Given the uncertainty associated with border molecules, we did not include them in our analysis.

**Figure 5.**
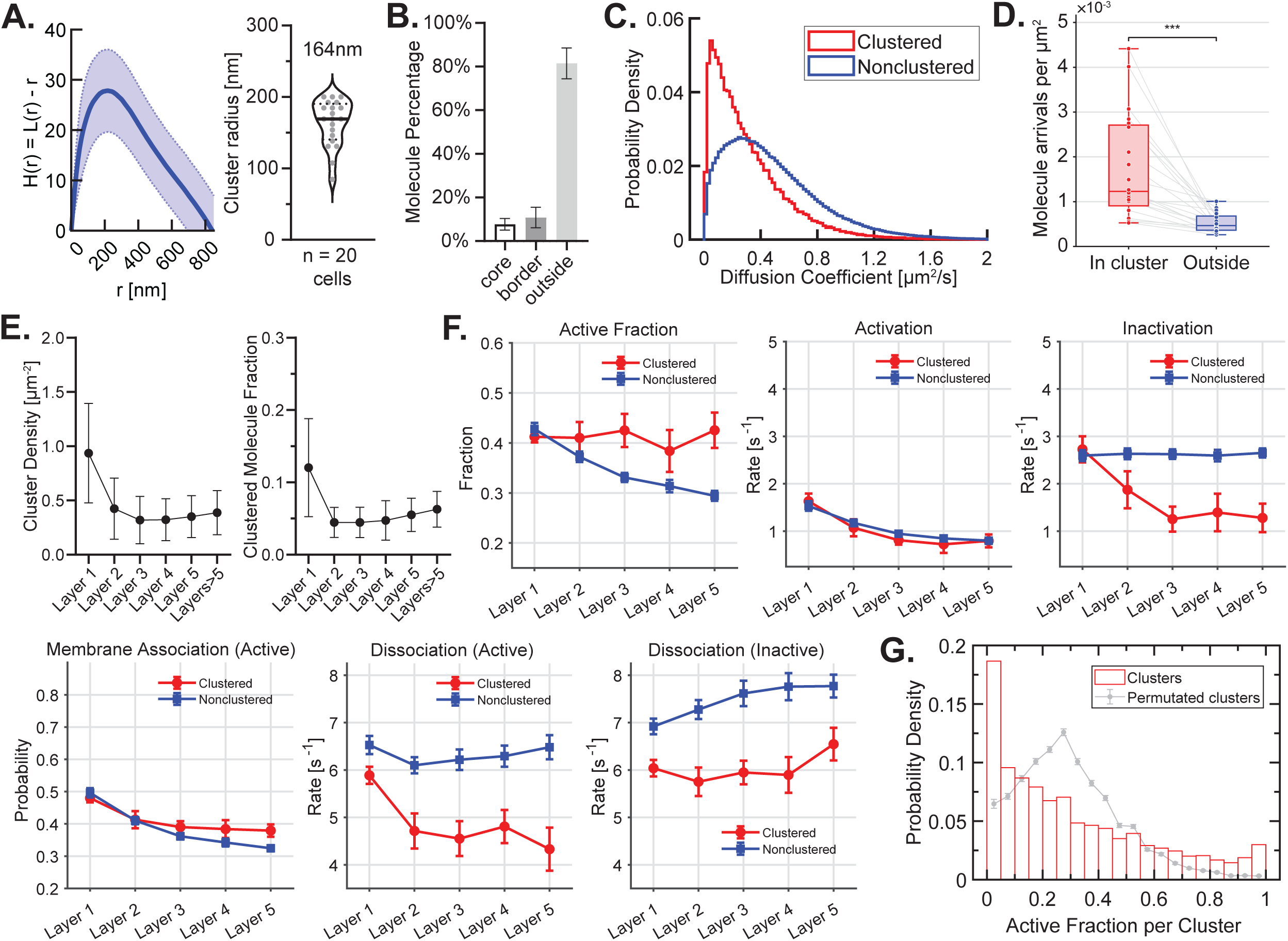
Analysis of Cdc42 nanoclusters. (A) Resulting H(r) function when all cells are combined (left panel) and estimated cluster radius for each cell (right panel). (B) Percentage of each molecule type. (C) Probabilities for the estimated diffusion coefficient for clustered and non-clustered molecules. (D) Number of molecular arrivals in 20 seconds per μm^2^ for clustered and non-clustered molecules. (E) Cluster density (left panel) and fraction in clustered molecules (right panel) in each layer. (F) Comparison of spatial rate profiles for clustered vs non-clustered molecules. (G) Probability density for the fraction of active molecules per cluster. The red bars are the experimental results and the gray data points are the results of a permutation test.

Core molecules comprised 8% of total Cdc42 (Fig 5B) and, consistent with incorporation in clusters, showed slower diffusion than nonclustered molecules (Fig 5C). The cluster density and fraction of clustered molecules was two-fold higher near the cell edge than in interior layers (Fig. 5E). The active fraction of clustered molecules was maintained throughout the layers, unlike non-clustered molecules where the active fraction decreased with distance from the edge (Fig. 5F). Several observations indicated that the clusters could function as localized sources of activated Cdc42 to produce activation gradients, or could be ‘hot spots’ where activated Cdc42 interacted with downstream molecules (*26, 27*).

Mechanisms for maintaining activation within clusters were suggested by our measurements. The activation rate did not differ for clustered versus non-clustered molecules, but the deactivation of clusters decreased by 60% over the first 3 layers, while deactivation of non-clustered molecules remained constant. Clusters had more molecule recruitment per unit area and lower rates of dissociation than did non-clustered molecules (Fig. 5D), for both active and inactive molecules. Interestingly, the fraction of clustered molecules that were active appeared skewed toward low and high values (Fig. 5G). To test if this observation was statistically significant, we performed a permutation test in which the label for clustered versus non-clustered tracks was randomly permuted (Fig. 5G, gray circles, and Fig. S4C); This clearly showed that the skewed distribution was not explained by random chance. We observed that clusters with dominantly inactive Cdc42 localizations (<10%) are less populated near cell edge, whereas other clusters are uniformly distributed across the cells (Fig. S4D). Together this data suggests that Cdc42 in clusters undergoes an organized lifecycle or that activation percentage is spatially controlled.

Our SPARC biosensors revealed conformational changes of individual proteins in living cells. They exemplify an approach to biosensor dyes that takes advantage of the bright signals and direct excitation of environment sensing dyes, making single molecule visualization and quantification possible. This capability revealed rates of conformational change in real time with sub-diffraction resolution, and enabled quantitation of rates for different steps in GTPase activation and membrane translocation. The primary driver of Cdc42 activity gradients driving cell protrusion was translocation of activated molecules to the membrane. We also observed a gradient of activation rates, but our computational model revealed this to play a secondary role. Analysis of single molecule tracks revealed clusters of Cdc42 molecules which were held at a high state of activity, unlike unclustered molecules, and with cluster density decreasing with distance from the edge. This provided a mechanism for localizing Cdc42 activity, providing hot spots for interaction with downstream effectors. Together these observations point to the importance of upstream molecules that control membrane translocation, indicate that activation occurs before membrane insertion or within 20 milliseconds of insertion, and reveal a role for Cdc42 clustering in maintenance and positioning of activity gradients.

## Methods

### Cell culture

Mouse embryonic fibroblasts (MEF, Clontech) and HeLa cells were maintained in 10% CO_2_ at 37 °C in culture medium composed of Dulbecco’s modified Eagle’s medium (DMEM, Cellgro) with 10% fetal bovine serum (HyClone, Thermo Scientific) and 2 mM GlutaMax (Gibco, Life Technologies).

The biosensor constructs were inserted into a Tet-Off inducible PiggyBac system and stable lines were produced in Tet-Off MEF cells (*28*). The stable MEF cells were cultured in DMEM and 10% FBS supplemented with 20 ng/mL doxycycline to suppress biosensor expression.

### Transfection, Dye Labeling, and Fluorescence Measurements

HeLa cells were plated into 6- or 12-well plates at a density of 40,000 cells or 20,000 cells per well, respectively. The following day, media was exchanged with fresh media containing 100 nM Nile Red SNAP (2 mL/well for 6-well dish, 1 mL/well for 12-well dish), and the cells were transfected with plasmid DNA using Fugene 6 (0.5 µg DNA/well of a 6-well dish, 0.25 µg DNA/well of a 12-well dish, 3:1 µL reagent/µg DNA). After another overnight incubation, the media was exchanged with dye-free media (1 or 2 mL/well) and cells were allowed to incubate for 1 hour to remove excess dye. The cells were then washed with PBS, trypsinized (200 µL/well of a 6-well plate, 150 µL/well of a 12-well plate), diluted with 4x medium, and each well was transferred into a separate Eppendorf tube. Cells were pelleted by centrifuging at 600 rcf for 5 minutes, the media was carefully aspirated off, and the cells were resuspended into PBS (500 mL if from 6-well plate, 400 mL if from 12-well plate). At this point, the emission spectrum of Nile Red was directly measured in the suspended cells. To measure Nile Red fluorescence using recombinant SNAP proteins, 500 nM dye in PBS containing 1 mM DTT was combined with a 5-fold excess of protein and was allowed to incubate at room temperature.

For single-molecule imaging, we achieved the appropriate density of labeled SPARC by tuning the biosensor expression and the concentration of dye used during labeling. To minimize signal from unattached Nile Red the dye concentration used for labeling and the washing procedure were optimized, leading to the protocol described below. If stable cell line was used for imaging, doxycycline was removed by transferring cells to new 6-well plates after wash. Cells were cultured in DMEM supplemented with 10% FBS and 200ng/mL doxycycline for 24 hours, then the cells were incubated with 200 nM Nile Red dye for another 12 hours. Dye solution in DMSO was added to cells in 1:1000 dilution. If transfection was used for imaging, HeLa cells were cultured in Phenol Red free DMEM plus 10% FBS for 24 hours. Then cells were plated into 6-well plates and transfected with SPARC constructs containing CMVd1 promoter using TransIT-X2 (Mirus Bio). 20 nM Nile Red was mixed into the transfection cocktail for simultaneous labeling for 24 hours. After biosensor expression and labeling, cells were incubated in dye-free FluoroBrite DMEM (Gibco) plus FBS and washed for at least 30 mins three times. Then cells were transferred into new dishes or coverslips and allowed to recover overnight before imaging. It is also recommended that imaging of positive and negative controls be performed for system calibration.

### Biosensor imaging

Biosensor imaging was performed on a home-built total internal reflection fluorescence microscope, equipped with four solid-state lasers (Coherent OBIS 405 nm, 488 nm, 561 nm and 647 nm). A four-band dichroic mirror (DM: Di01-R405/488/561/635, Semrock) was used for multi-color imaging. Fluorescence images were collected using a 60 X TIRF objective (UAPON 60XOTIRF, NA 1.45, Olympus) and split by a dichroic mirror (Semrock, FF624-Di01) through a dual-camera system (Hamamatsu, W-VIEW GEMINI-2C). The images were projected onto two metal-oxide-semiconductor (sCMOS) cameras (Photometric, 95B Prime). The laser power was set to 1 mW and the exposure time was set to 100 ms with 3 seconds interval between time points. After background subtraction and registration, the shorter wavelength image was divided by the longer wavelength image to produce a ratio image reflecting Cdc42 activity.

MEFs labeled with the dye JF646 were imaged using an inverted IX81 epifluorescence microscope (Olympus) with a 40x 1.3NA Silicon oil objective (UPLSAPO40XS, Olympus) and a ORCA-Flash4.0 V2+ sCMOS camera (Hamamatsu). Images were collected every 10 sec for a total of 15 min using FF405/496/593/649-Di01 as a dichroic (Semrock), HQ630/40X and HQ470/40X (Chroma) as excitation filters and ET705/72 and HQ525/50M (Chroma) as emission filters (for JF646 and GFP respectively).

Ratiometric image processing was as done following previously described procedures (*29*). Briefly, the raw images were passed through a processing pipeline including shade correction, baseline subtraction, masking, registration, and photo-bleach correction when needed, then ratiometric images were obtained by dividing two images. The range of the color bar was set to the 5th and 95th percentiles of the ratio images.

### Fluorescence lifetime imaging

FLIM was carried out on a Lecia TCS SP8 X confocal microscope equipped with a pulsed White Light Laser (WLL2, 470-670 nm), using a 40x/1.30 Plan Apochromatic (oil) WD = 0.24 mm objective. Excitation was at 561 nm and emission was collected at 570-670 nm. Leica Application Suite X (LAS X) software platform was used for image processing and phasor analysis.

### Single-molecule image analysis

#### Single Molecule Tracking

During live-cell imaging, photons were filtered into 2 channels based on their wavelengths. After dark current subtraction, a baseline of 800 intensity units was added to the dark current corrected images to avoid negative intensities and preserve the shape of the point spread function. Individual molecules were tracked in the two channels separately using the software package u-track (*30*). For particle detection, sigma of the point spread function (psfSigma) was set to 1.4 and alpha for detection significance (alphaLocMax) was set to 0.18. Multiple Gaussian fitting was enabled to estimate the positions of molecules whose intensities overlapped. To connect measurements from the two channels, pairwise distances of all particles detected in the two channels were measured. To be considered the same molecule, pairs had to be within 4.9 pixels of each other. If more than two molecules satisfied this condition, the closest two were considered the same molecule. The centroid of paired detections was used as the molecule’s position. During tracking, a maximum time gap of 1 frame was allowed to account for blinking, and merging/splitting was enabled. Tracks shorter than 2 frames were discarded.

#### Local Background Subtraction

For both channels, fluorescence intensity of the molecules was calculated as the sum of the intensities in a 3-pixel radius disk (Fig. S2D). We considered a concentric ring of width 1 pixel around the disk the background intensity (Fig. S2D). The intensities in the ring were summed and scaled to account for the larger area of the ring. This number was then subtracted from total intensity inside the disk.

#### Emission Spectra Scaling

We scaled the measured fluorescent intensities to account for cell-to-cell variability before using them in the hidden Markov model (HMM). WT, negative control and positive control cells were scaled separately. The first step in the scaling process was to manually choose a single dividing line to separate the intensity measurements for all cells (Fig. S2E, top row, red line). For each cell, the intensities above the line were used to find the mode of the active population by projecting these intensities onto the y-axis, and the intensities below the line were used to find the mode of the inactive population by projecting the intensities onto the x-axis. The data for each cell was then scaled by dividing by the modes (x/xmode, y/ymode) so that both modes lay at 1 in the standardized distribution (Fig S2E, bottom row).

#### Hidden Markov Model to Infer Cdc42 States and Kinetic Rates

We used a hidden Markov model (HMM) with the hmmlearn Python package to infer the state of individual Cdc42 molecule at each observed time point and estimate the transition rates in the GTPase cycle. HMMs are used to infer system states that are not directly observable but functionally related to measurable quantities. In our case, the states of individual Cdc42 molecules (active or inactive) are not observable but inferred from fluorescent time series measurements. To be considered in our analysis duration of single molecule tracks had to be at least 5 frames or 100 ms.

The HMM relies on three sets of probabilities: the starting probabilities, the emission probability distributions, and the transition probabilities. The starting probabilities refer to the state that a Cdc42 molecule arrives at the cell membrane (active or inactive). The emission probability distributions are based on the distribution of observable signal emitted by each state. We used the ratio of the fluorescent signal in the short and long wavelength channels as the emission probabilities. Active molecules exhibit higher ratios than inactive molecules due to the spectral shift to the shorter wavelengths. The ratio of two Gaussian distributed random variables is lognormally distributed (Fig. S2C). Therefore, we chose the GaussianHMM model in hmmlearn and supplied the natural logarithm of channel ratios as input. The third set of probabilities the HMM utilizes are the transition probabilities for changes in the state of a Cdc42 molecule between successive time frames. These transition probabilities are related to the reaction rates in the GTPase cycle.

Given time series of observed signals, an iterative expectation-maximization algorithm known as the Baum-Welch algorithm can be used to estimate all model parameters and infer the most likely states at each time point. However, it is not guaranteed that any given dataset will contain enough instances of active and inactive molecules to infer the emission probability distributions. To avoid inaccuracies between datasets, we first ran the hmmlearn package with a pool of tracks from standardized negative control, positive control and wild type cells. This provided us with Gaussian distribution parameters that best approximated the emission probability distributions of the active and inactive states (Fig. S2F). To determine the spatial dependence of the transition rates, we reran the HMM on tracks selected from different regions of the cell using the emission probabilities inferred from the pooled run.

#### Kinetic Rate Estimation

The transition probability matrix P and corresponding rate matrix Q for the three-state system (active A, inactive I, and absorbing Ø for dissociation from the membrane) have the following forms:

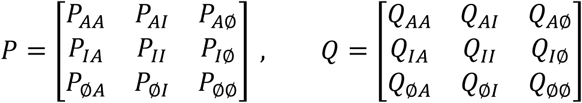

P and Q are related by the function:

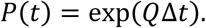

where Δt is the time interval between measurements.

The transition probability matrix was calculated using the states inferred with hmmlearn and the transition rates were calculated by taking the matrix logarithm of P in MATLAB.

### Nanocluster analysis

Ripley’s K function (*31*) was applied to Cdc42 track centroids using customized MATLAB scripts. The function describes the difference between pairwise particle distances and the value expected from uniformity. Positive values in L(r)-r function indicates clustering against homogenous distribution. We further estimate the radius of the clusters using local maxima position of the K function divided by empirical coefficient 1.3 as described in Ruan et al. (*32*).

We identified Cdc42 clusters by applying Density-based spatial clustering of applications with noise (DBSCAN) (*25*) to track centroids using customized MATLAB scripts. DBSCAN depends on two variables: the total number of points (*N*) within a certain distance (*d*). We required that a molecule must be at least d = 140nm of any given molecule to be considered as a cluster-forming molecule and a cluster must contain at least N = 15 molecules. DBSCAN reports two groups of molecules in clusters: core points and border points. Core points satisfy the aforementioned criteria. Border points are within d = 140nm of core points but do not have N = 15 molecules in their local neighborhood. Activation kinetics of border molecules are similar to non-cluster molecules. Therefore, we only use core molecules from DBSCAN in subsequent analyses.

Since relatively few molecules are present in clusters, tracks across cells were pooled together for parameter estimation. The standard error of the parameters across cells was estimated by bootstrapping with 800 bootstraps with replacement.

The permutation test was done using MATLAB. For all tracks in one cell, track indices were randomly shuffled. The position of each track was translated so that its new center matched the center of the original (unpermuted) track. The active fraction of locations in each cluster was reevaluated. A single permutation round used results from 20 cells and was repeated 1000 times.

### PDE Simulations

The reaction-diffusion system for Cdc42 activity was coded in MATLAB and solved with the ode15s solver using a second-order central difference for the spatial second derivative. The model consists of the following three variables: *C*(*t*) is the concentration of the well-mixed cytosolic pool of Cdc42; *I*(*x*, *t*) is the concentration of inactive, membrane-bound Cdc42; *A*(*x*, *t*) is the concentration of active, membrane-bound Cdc42. The spatial domain: *x* ∈ [0, *L*], where the cell edge is located at x = 0. The model equations are:

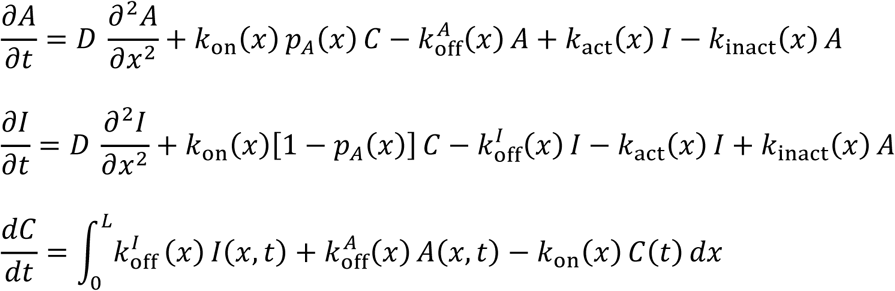

with no flux boundary conditions:

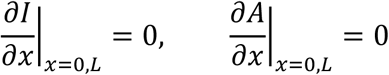

Reaction rates at each spatial grid point were obtained by interpolating between the experimentally measured reaction rates with the makima method. The spatial profile of total Cdc42 concentration was used to scale the association rate. Reaction rates were set to the mean of the corresponding spatially dependent rate to test the impact of the gradient.

### Biosensor engineering

To develop SPARC Cdc42, we first fine-tuned the linkers connecting CBD, SNAPtag, and Cdc42 by comparing biosensors containing a constitutively active Cdc42 (Q61L) with the same constitutively active biosensor containing additional mutations in CBD to reduce Cdc42-CBD binding (H246D and H249D). We assessed differences in emission maxima and the ratio of areas under the two emission bands used for live cell experiments (Fig. S1B). Fluorescence spectra were obtained from suspensions of HeLa cells expressing biosensor protein, after incubation with dye and washing. When we tested flexible and soluble Gly-Ser-Gly (GSG) linkers, we found that the difference between the active and inactive state was improved as the linkers were shortened (Fig. S1B, GSG linker), likely because this restricted the orientation of the dye that interacted with the CBD-Cdc42 complex, and possibly also because shorter linkers reduced the entropy loss upon dye binding. Ultimately, the greatest difference was obtained by using no linkers. To further decrease the spacing between biosensor components, we tested truncations of the CBD and SNAP proteins. A large portion of the CBD C-terminus is not involved in the interaction with Cdc42 (32), so we removed the last 25 amino acids of CBD in 5 amino acid increments (Fig. S1B, CBD trunc.). The biosensor’s dynamic range increased when up to 15 amino acids were removed (CBD-15). Removing more was not effective, likely because this prevented CBD from reaching Cdc42. We truncated the N- and C-termini of the SNAP-tag, and also removed the restriction sites (RS) used at each terminus for cloning (Fig. S1B, RS detel. & SNAP trunc.). Removing 6 amino acids greatly reduced dye attachment, as this portion of SNAP contains a Cys that binds Zn2+ to stabilize SNAP. Interestingly, some modifications caused a red shift of the dye emission. The greatest improvement was achieved by combining truncation of 2 amino acids with removal of the C-terminal restriction site. Combining all these optimizations produced a 20 nm emission shift and 2.5-fold change in area ratio upon activation (Fig. S1B).

To further improve SPARC Cdc42, we experimented with hybridization of two Cdc42 binding peptides, CBD and the GTPase-binding domain from PAK 1 (PBD). PBD recognizes the active conformation of both Rac1 and Cdc42 and has been used in FRET-based Cdc42 biosensors. Aligned CBD and PBD sequences were divided into six segments (A-E) based on crystal structures (Fig. S1A). We generated the chimeric peptides by swapping these short fragments between CBD and PBD. Surprisingly, replacing CBD with PBD in the Cdc42 biosensor design abolished almost all the response (Fig. S1C). To understand which portions of CBD and PBD were important for binding, we substituted PBD fragments into the CBD affinity reagent. Swapping PBD A-C completely abolished response, while PBD D-F increased response (Fig. S1C top left panel). Swapping CBD B-F with PBD produced little or no improvement, while substituting in A led to a response as robust as native CBD (Fig. S1C top right panel), suggesting CBD A is critical for sensor response. Simply replacing region F of CBD with F from PBD increased the activation-induced shift to 32 nm (Fig. 1B left panel).

To test whether the design was generalizable, we tested an analogous SPARC biosensor for the small GTPase Rac1. SPARC Rac1 response was increased by placing PBD D and F into CBD, and by inserting CBD A with PBD (Fig. S1C bottom panel). These studies showed that regions A-C were critical for dye response, so we studied the N-terminus in more detail. We pursued a better Rac1 biosensor by focusing on fine tuning the A-C regions. Seven subregions (M1-M7) of CBD were substituted with PBD fragments to make Rac1 biosensors. Separation of subregions was based on residual hydrophobicity scores (Fig. S1D). Region M1 and M5 produced the largest activation-dependent change (Fig. S1E right panel). When combined with the previous PBD D&F swap, we obtain a 26 nm shift. This was the fully optimize biosensor we named SPARC Rac1 (Fig. S1E right panel). Interestingly, M1-M7 subregion substitution did not substantially improve SPARC Cdc42 (Fig. S1E left panel).

### Synthesis of Nile Red SNAP

See Supplement Note for detailed synthesis procedures of Nile Red SNAP.

---

**Supplemental Figure 1.**
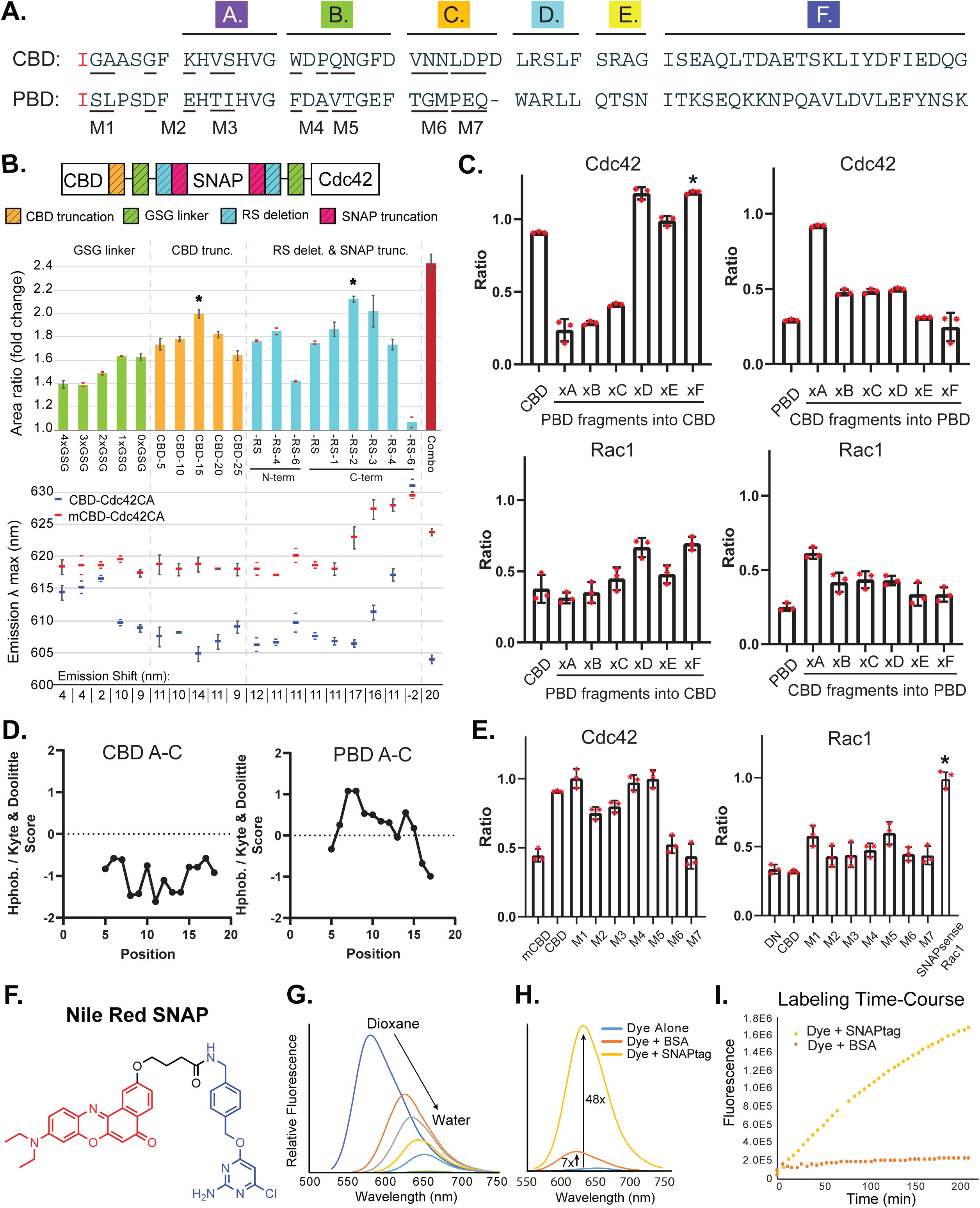
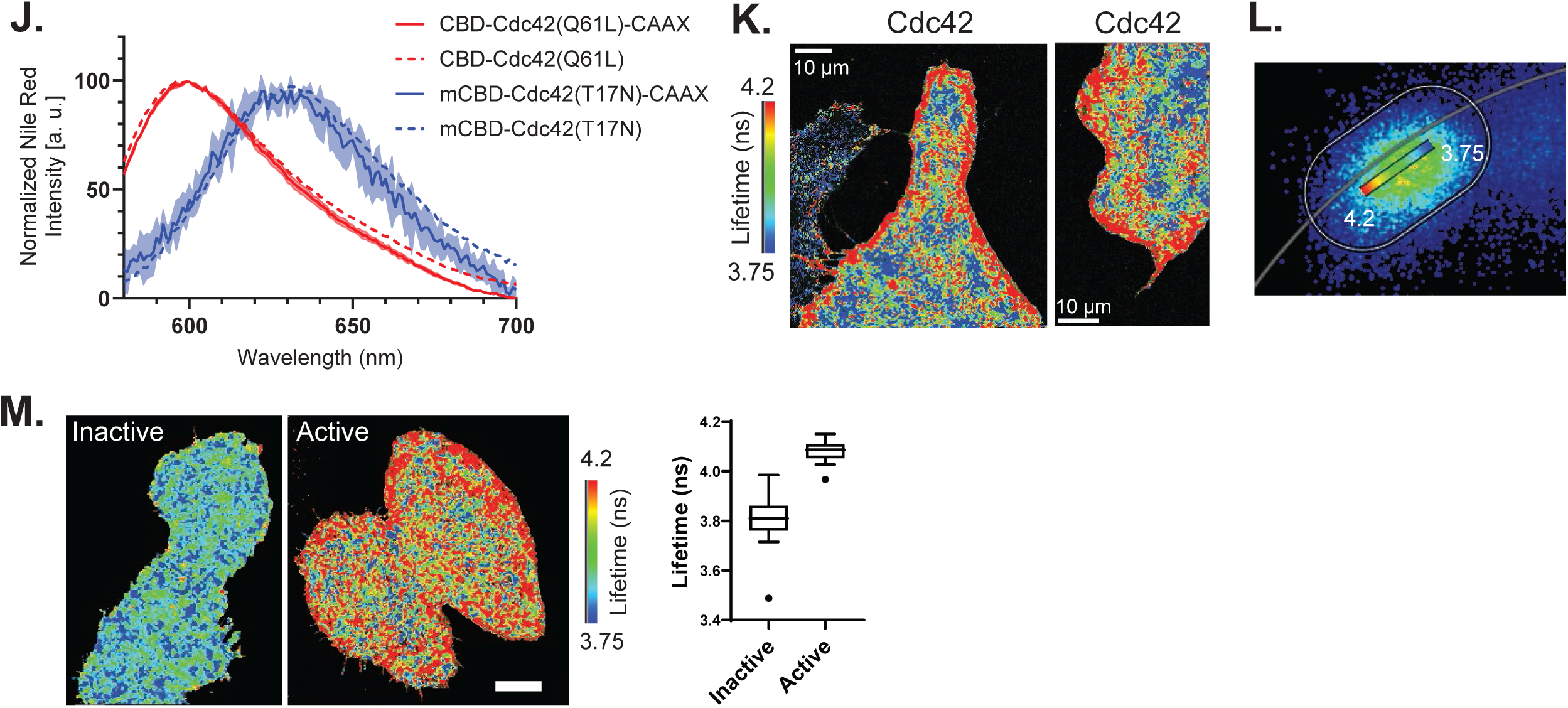
Biosensor development (A) Sequences of the CBD and PBD domains showing segments swapped to produce affinity reagents with optimum brightness and fluorescence change. CBD is derived from Wiskott Aldrich Syndrome protein (WASp) and PBD from Pak1. (B) The ratio of areas under the emission curve (AUC) for emission below and above 624 nm, and the change in emission maxima, induced by varying linkers, truncating CBD, and deleting restriction sites. Combo = optimal combination including all starred modifications. Data is plotted as the mean ± s.d.; n = 3 biologically independent samples when error bars are shown, otherwise n = 2. Individual ratio values above and below the mean. (C) SPARC Cdc42 and Rac1 optimization by combining CBD and PBD domains. The emission ratios of activated (Q61L) mutants are shown. Ratio changes produced by substituting PBD segments into CBD (left), or CBD into PBD (right). Segments shown in Fig. S1A. Final SPARC Cdc42 biosensor is marked with *. (D) Point mutations M1-M7 were selected and tested based on hydrophobicity scores. (E) Point mutations (see Fig. S1A) produced no significant improvement in SPARC Cdc42. SPARC Rac1 incorporated point mutations M1 and M5 from PBD into CBD, and replaced the D and F fragments of CBD with D and F fragments of PBD. Final SPARC Rac1 biosensor is marked with *. (F) The SNAP-reactive Nile Red derivative (NR-SNAP) used in the biosensor, consisting of Nile Red (red), a linker (black), and a SNAP-reactive group (blue). (G) Emission of equimolar NR-SNAP in mixtures of dioxane and water, showing hypsochromic shift and decrease in intensity as solvent polarity increases (100% dioxane to 100% water, in 20% increments.) (**H, I**) Response of NR-SNAP when linked to SNAP-tag. (H) Normalized spectra show the emission changes produced by labelling SNAP-tag versus non-specific interaction with BSA. (I) Time course of fluorescence increases during SNAP-tag labelling versus interaction with BSA. (J) Replacing the C-terminal CAAX sequence of SPARC Cdc42 with that of KRas for constitutive membrane localization does not change the emission spectrum, showing that the Nile Red spectral shift does not result simply from membrane translocation. (K) FLIM imaging of SPARC Cdc42 in a MEF cell shows activation at the edges of protruding regions. (L) Fluorescence lifetime phasor plot from HeLa cells expressing wildtype SPARC Cdc42. (M) FLIM imaging of Hela cells expressing the inactive and active Cdc42 biosensor mutants (Q61L, T17N) and overall lifetime measurements of single cells expressing the indicated constructs.

**Supplemental Figure 2.**
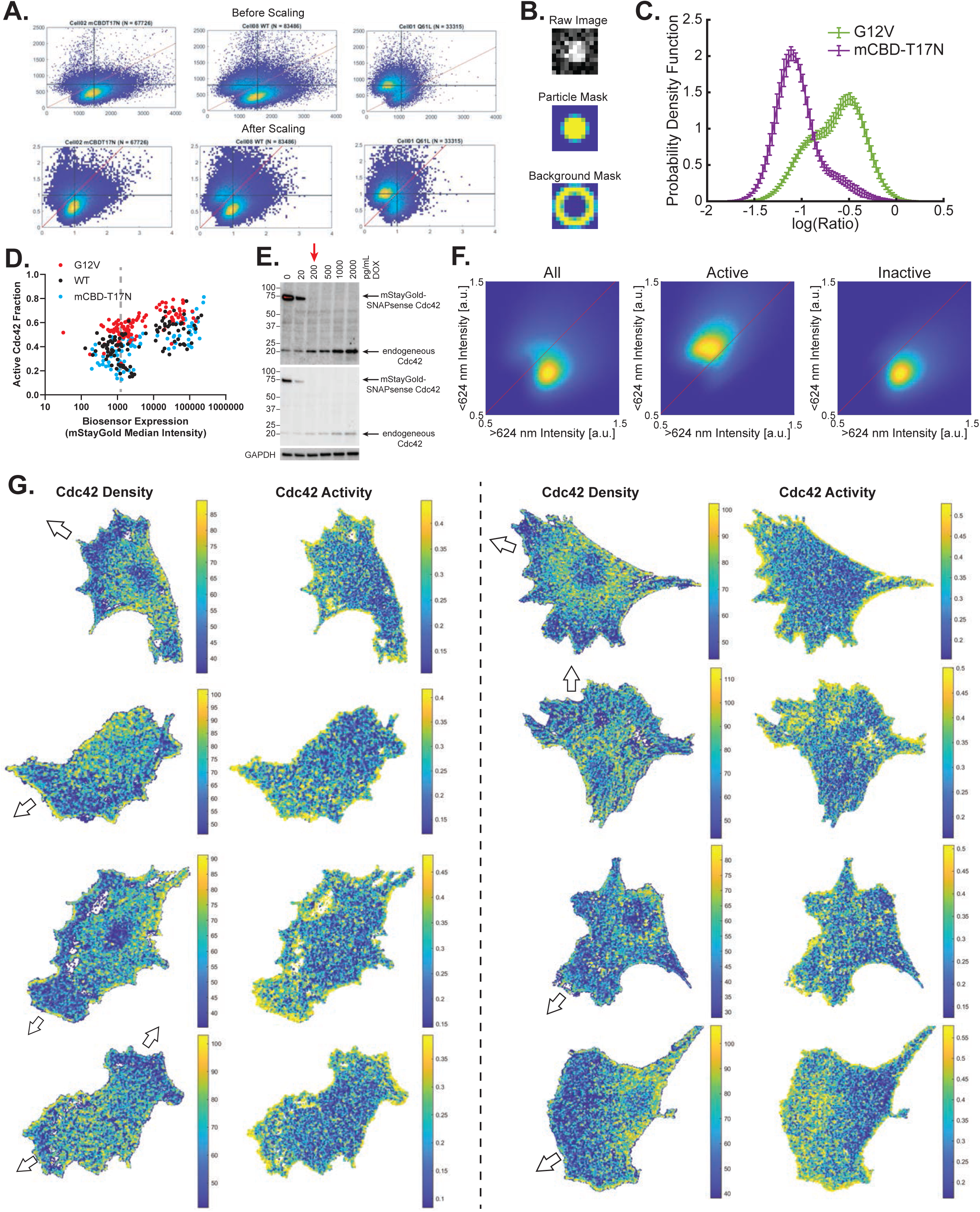
**Single-molecule ratio imaging using SPARC Cdc42.** (A) Single-molecule intensities are normalized based on the mode intensity in each wavelength channel to reduce variabilities between datasets. (B) Single-molecule intensity is extracted by applying a particle mask to the raw image. Background subtraction was performed using the background mask as indicated. (C) Histogram of single-molecule ratios from constructs shown in Fig. 2B. (D) Cdc42 activation increases with biosensor expression. Note logarithmic scale. Endogenous Cdc42 level is indicated by the gray dashed line. (E) Biosensor expression under the control of doxycycline. Doxycycline concentrations were 0 pg/ml, 20 pg/ml, 200 pg/ml, 500 pg/ml, 1 ng/ml, and 2 ng/ml. The gel image is shown at two contrast settings. (F) 2D histogram of single-molecule intensities for all molecules, active Cdc42, and inactive Cdc42 based on HMM training. (G) Density versus activity of SPARC Cdc42 tracks in moving cells. Each point marks the centroid of a single track. In the density map, color indicates the number of molecules within 500nm. In the activity map, it shows the fraction of active molecules within 500 nm. Scale bar: 10 μm.

**Supplemental Figure 3.**
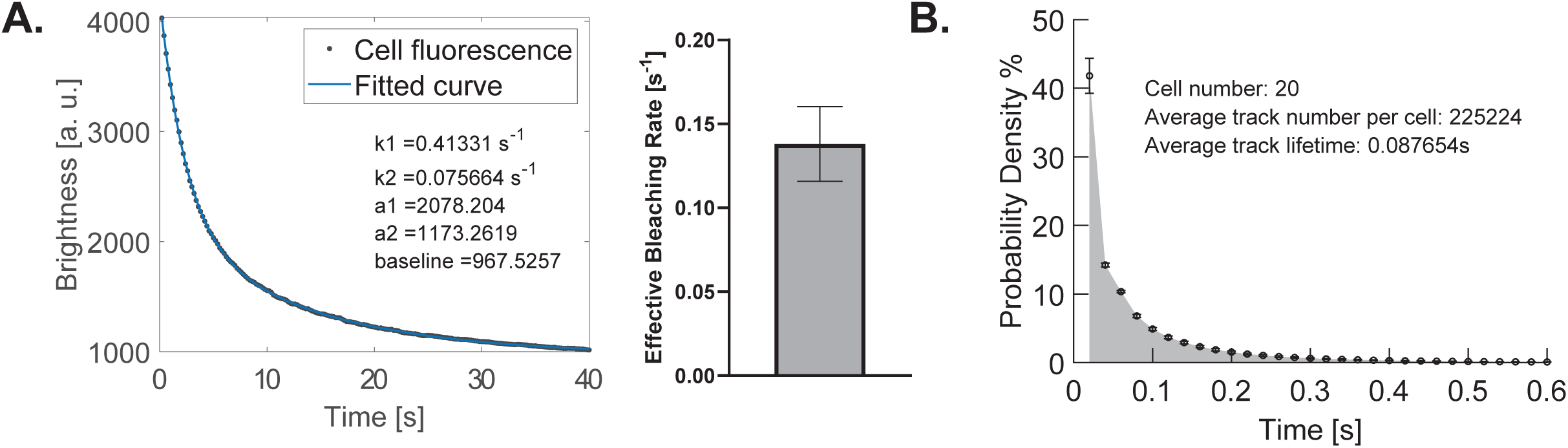
Single-molecule kinetics of Cdc42 in motile MEF cells. (A) Bleaching rate calculation for SPARC Cdc42. We fit a double exponential decay function to the biosensor brightness over time (left). The effective bleaching rate is the harmonic mean of the two rates from the fitting (right, cell number = 28). (B) Track lifetime distribution for SPARC Cdc42. Mean track lifetime is 0.087654s from 225224 tracks summed over 20 cells.

**Supplemental Figure 4.**
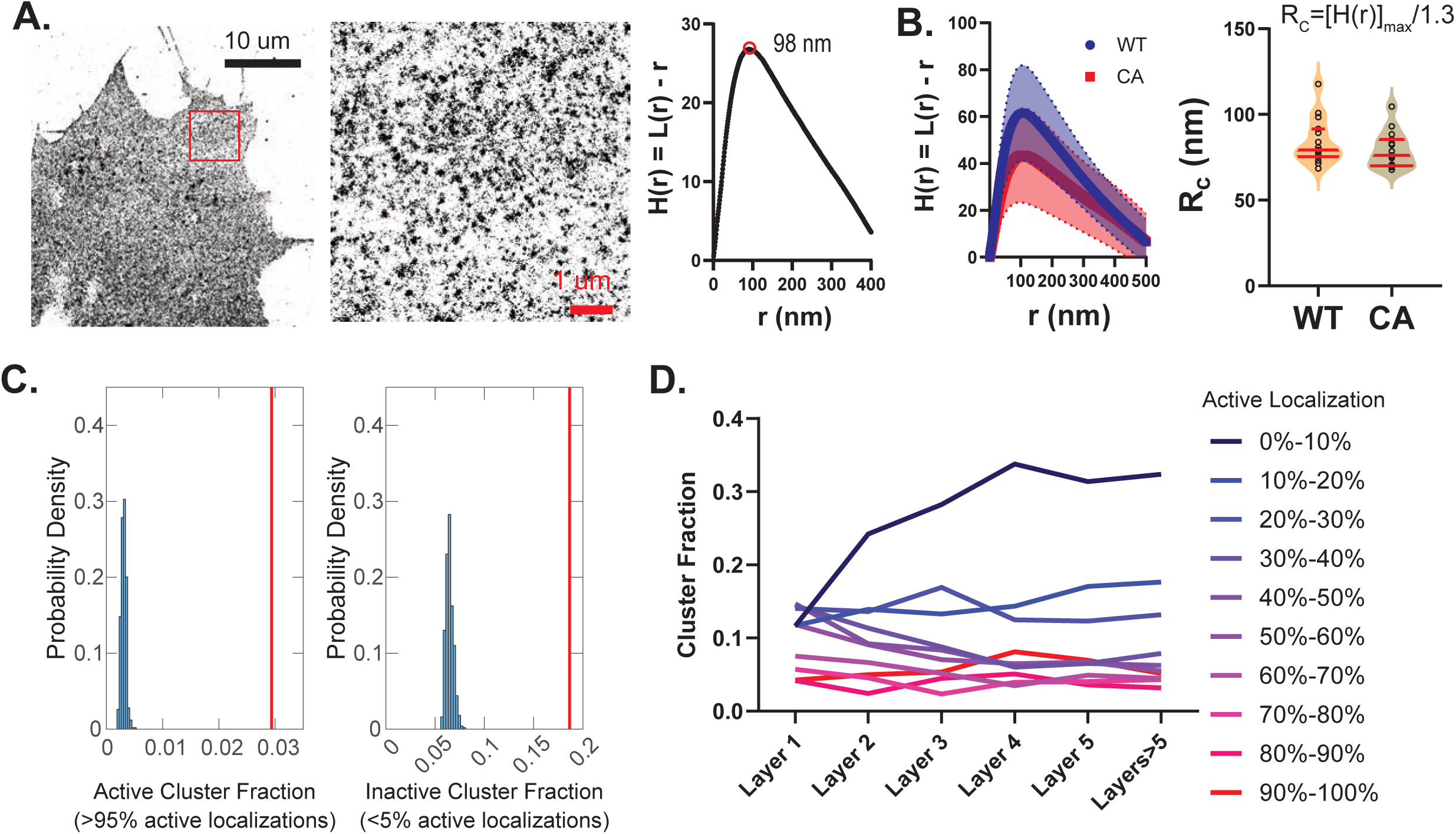
Cdc42 form clusters on the plasma membrane. (A) PALM reconstruction of a fixed MEF cell expressing Cdc42-PAmCherry revealed that Cdc42 was organized in clusters on the PM. Shown in the middle is a zoomed-in view of the region marked red in the left panel. Ripley’s K function on the right showed the average cluster size to be ∼100 nm. (B) Left: Ripley’s K function plot of 5 μm x 5 μm regions from cells expressing wildtype and constitutively active Cdc42. Note that higher peak values indicated that the clusters were more separated. Right: Quantification of the corrected cluster radius for wildtype and constitutively active Cdc42. WT: N= 15; CA: N=15. (C) Histogram of active and inactive cluster fractions from permutation test. Active clusters contain more than 95% active localizations and inactive clusters contain less than 5%. Biological active cluster fraction is ∼3% and inactive cluster fraction is ∼18%, significantly higher than that from permutation test. (D) Spatial variation of clusters with different activity levels. Clusters containing less than 10% active localizations were downregulated in regions near the cell edge. The rest of the clusters show even distribution across the cell.

**Supplemental Table 1.**
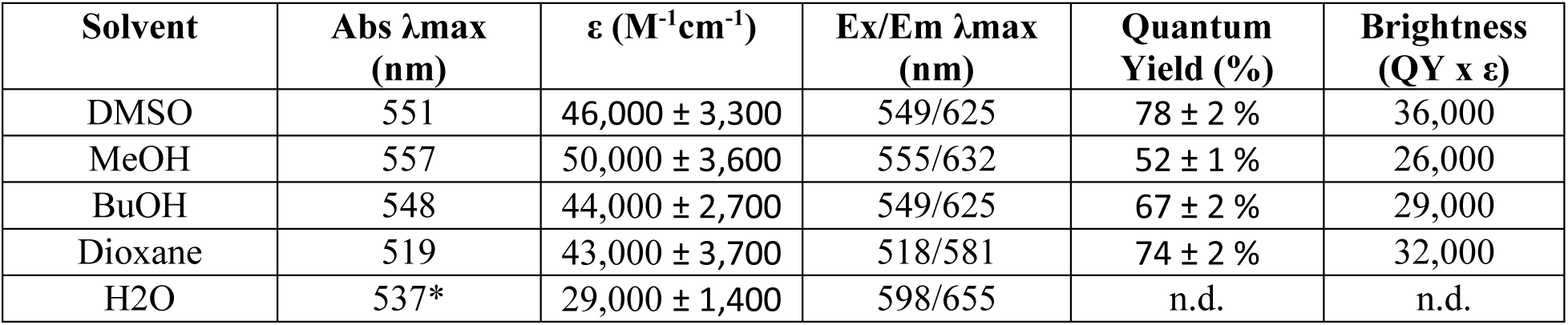
Photophysical properties of Nile Red SNAP.

| **Solvent** | **Abs λmax (nm)** | **ε (M<sup>-1</sup>cm<sup>-1</sup>)** | **Ex/Em λmax (nm)** | **Quantum Yield (%)** | **Brightness (QY x ε)** |
|---|---|---|---|---|---|
| DMSO | 551 | 46,000 ± 3,300 | 549/625 | 78 ± 2 % | 36,000 |
| MeOH | 557 | 50,000 ± 3,600 | 555/632 | 52 ± 1 % | 26,000 |
| BuOH | 548 | 44,000 ± 2,700 | 549/625 | 67 ± 2 % | 29,000 |
| Dioxane | 519 | 43,000 ± 3,700 | 518/581 | 74 ± 2 % | 32,000 |
| H2O | 537* | 29,000 ± 1,400 | 598/655 | n.d. | n.d. |

## Acknowledgements

We thank Luke Lavis (Janelia Research Campus) for providing Janelia Fluor dyes and Mauro Calabrese (University of North Carolina at Chapel Hill) for vectors used to generate stable cell lines. We gratefully acknowledge the National Institute of General Medical Sciences for financial support (F32GM120958 to NKP, R35GM127145 to TCE, and R35GM122596 to KMH).

