## Supplemental Note for "Tracking the conformational dynamics of single molecules in live cells; GTPase cycling in Cdc42 gradients and microclusters"

### Supplemental Info

#### General materials and methods

All reagents and solvents were purchased from commercial suppliers (TCI America, Acros, Sigma Aldrich, Fisher Scientific) and used without further purification. TLC chromatography was performed using TLC Silica gel 60G F254 25 Glassplates (EMD Millipore). UV-visible spectra were obtained with a Hewlett-Packard 8453 diode array spectrophotometer. Chromatography was carried out on an Isco Combiflash Companion apparatus. Emission and excitation spectra were obtained using a Horiba Fluorolog 3 spectrofluorometer at room temperature. Mass spectra were obtained on a ThermoScientific Q Exactive HF-X using direct infusion. NMR spectra were obtained on an Inova-400 spectrometer. Chemical shifts are reported using NMR reference peaks of 7.26 ($^1$H) and 77.23 ($^{13}$C) ppm for $CDCl_3$, and 2.50 ($^1$H) and 39.51 ($^{13}$C) ppm for DMSO-$d^6$.

#### Synthesis of Nile Red SNAP

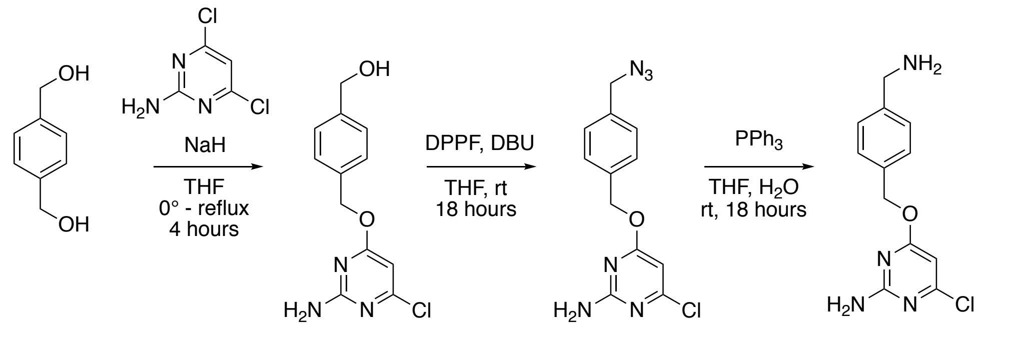


**Supplemental Scheme 1.** Synthesis of SNAP-$NH_2$.

**(4-(((2-amino-6-chloropyrimidin-4-yl)oxy)methyl)phenyl)methanol (1).** Benzene dimethanol (3.37 g, 24.4 mmol) and 4-Amino-2,6-dichloropyrimidine (1.0 g, 6.1 mmol) were dissolved into dry THF (11 mL) under argon and cooled to 0 °C. Sodium hydride (60% dispersion in mineral oil, 234 mg, 6.1 mmol) was added in one portion and the reaction was allowed to continue to stir at 0 °C for 10 minutes. Then the reaction was brought to reflux for four hours. The reaction was cooled to room temperature and quenched with methanol, then was dried onto silica gel. The product was purified on silica gel using a gradient of 0-10% methanol in DCM to yield a yellow solid (1.21 g, 80% yield).

$^{1}H$ NMR (400 MHz, DMSO-$d_6$) δ 7.39 (d, *J* = 8.1 Hz, 2H), 7.32 (d, *J* = 8.1 Hz, 2H), 7.10 (s, 2H), 6.14 (s, 1H), 5.30 (s, 2H), 5.18 (t, *J* = 5.7 Hz, 1H), 4.49 (d, *J* = 5.6 Hz, 2H). $^{13}C$ NMR (100 MHz, DMSO-$d_6$) δ 170.33, 162.79, 160.00, 142.53, 134.49, 128.17, 126.43, 94.42, 67.34, 62.61. HRMS (M + $H^+$): Expected: 266.0691, Found: 266.0681.

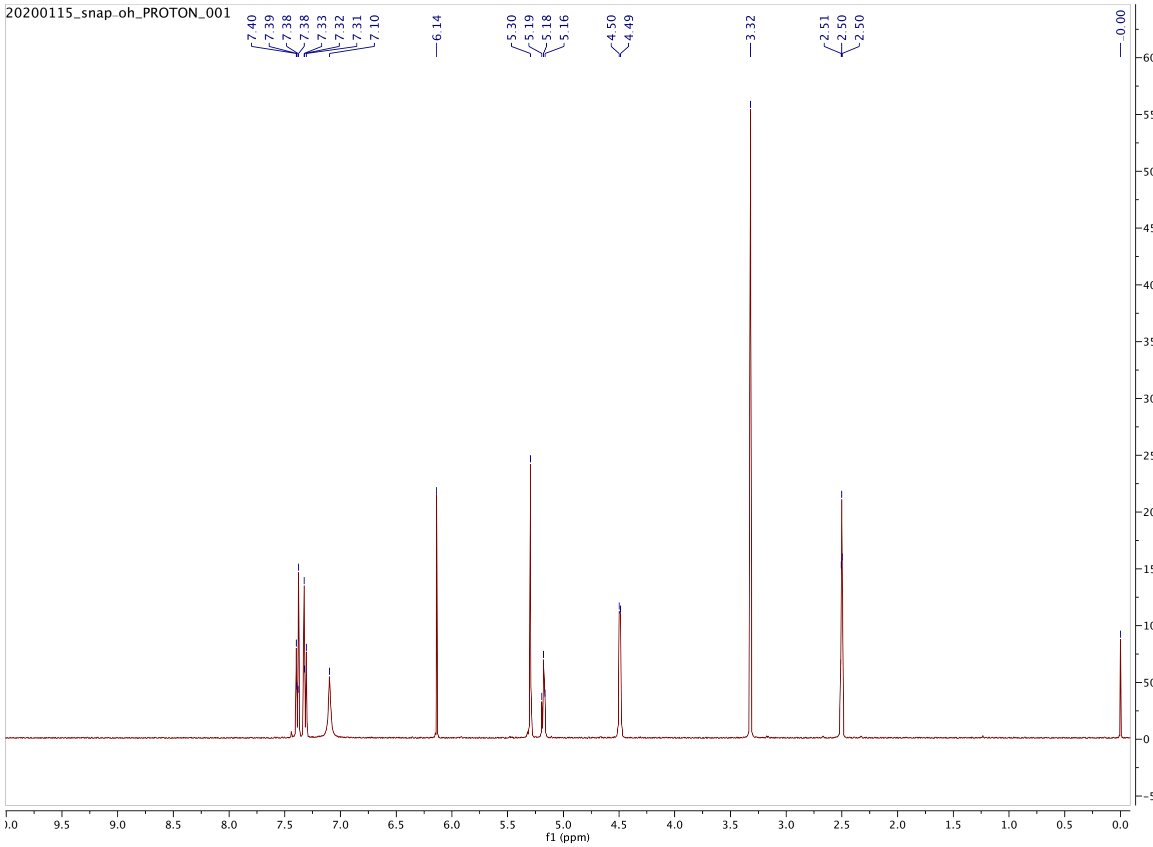

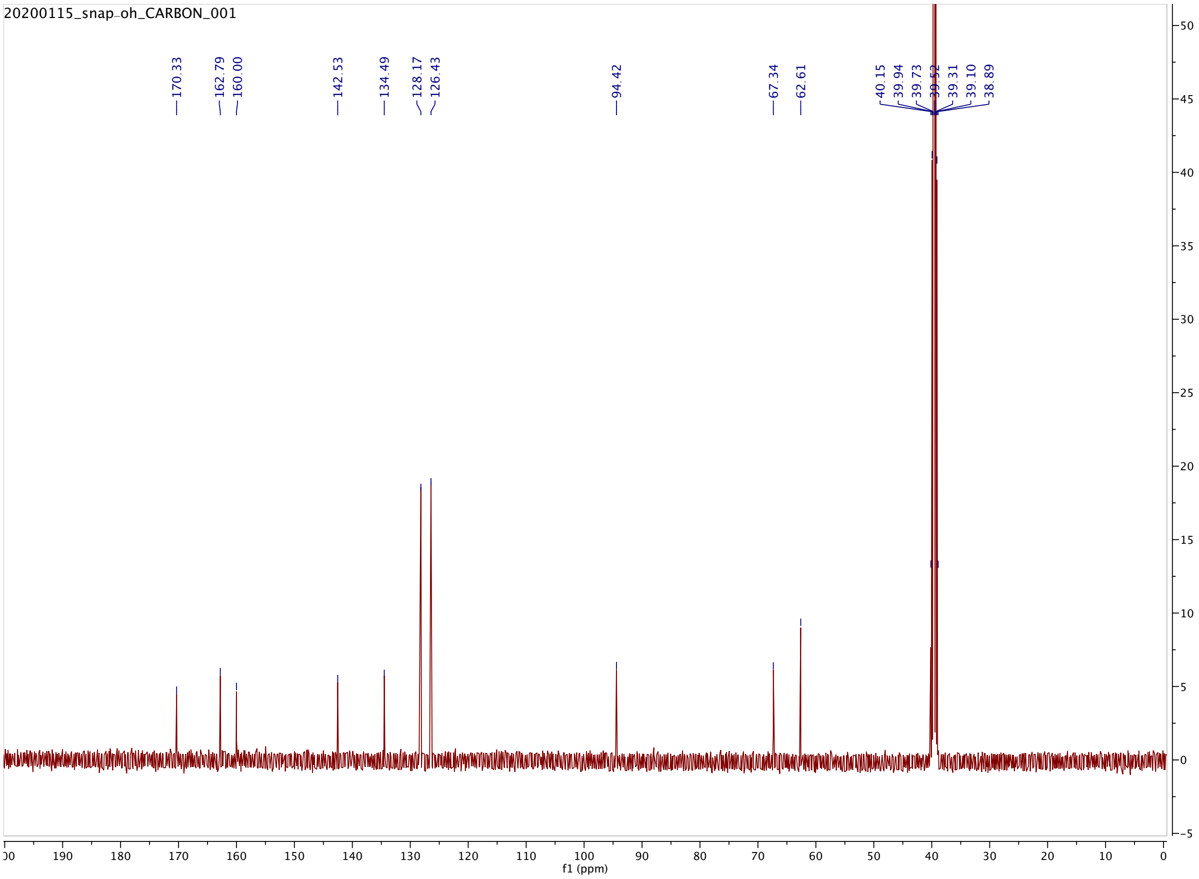

20200115_snap-oh_CARBON_001
170.33
162.79
160.00
142.53
134.49
128.17
126.43
94.42
67.34
62.61
40.15
39.94
39.73
39.52
39.31
39.10
38.89
f1 (ppm)

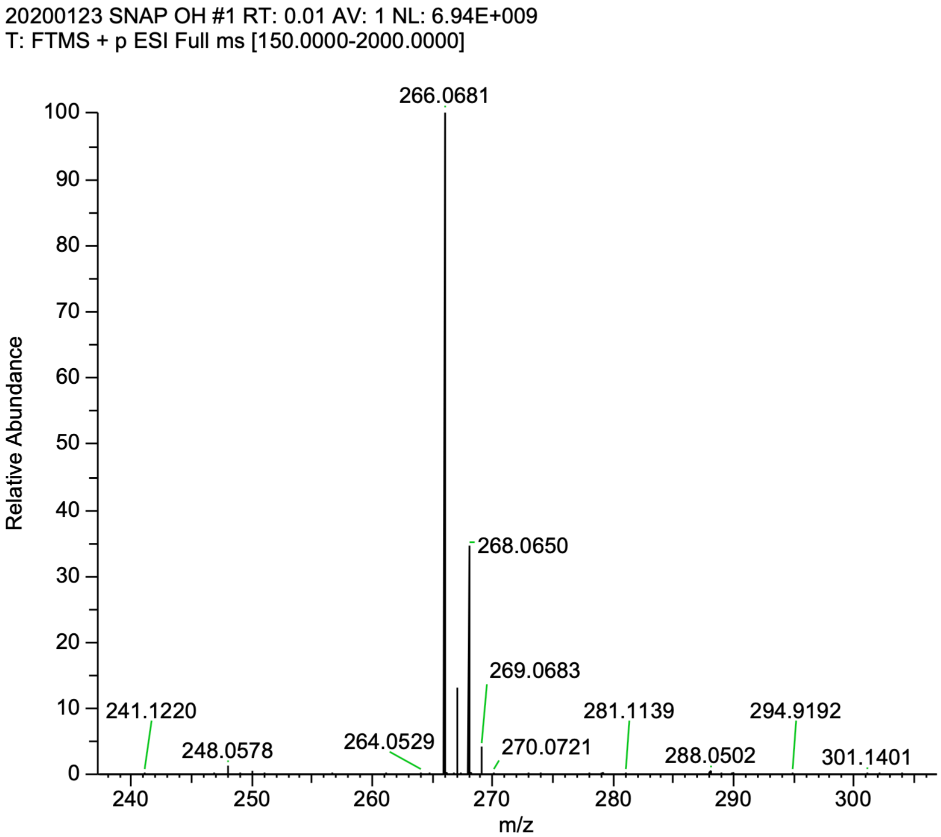


**4-((4-(azidomethyl)benzyl)oxy)-6-chloropyrimidin-2-amine (2). 1** (500 mg, 1.85 mmol) was dissolved into dry THF (38 mL) under argon, then DBU (0.414 mL, 2.78 mmol) and diphenylphosphoryl azide (DPPA, 0.476 mL, 2.21 mmol) were added. The reaction was allowed to stir at room temperature overnight, after which TLC showed complete conversion to product. The reaction solution was dried onto silica gel and the product purified with a gradient of 10-40% ethyl acetate in hexanes to yield a fluffy white solid (503 mg, 99% yield).

$^{1}$H NMR (400 MHz, DMSO-$d_6$) δ 7.47 (d, 2H), 7.39 (d, 2H), 7.11 (s, 2H), 6.16 (s, 1H), 5.33 (s, 2H), 4.45 (s, 2H). $^{13}$C NMR (100 MHz, DMSO-$d_6$) δ 170.28, 162.79, 160.04, 136.19, 135.46, 128.60, 128.52, 94.41, 67.06, 53.26. HRMS (M + H$^{+}$): Expected: 291.0756, Found: 291.0744.

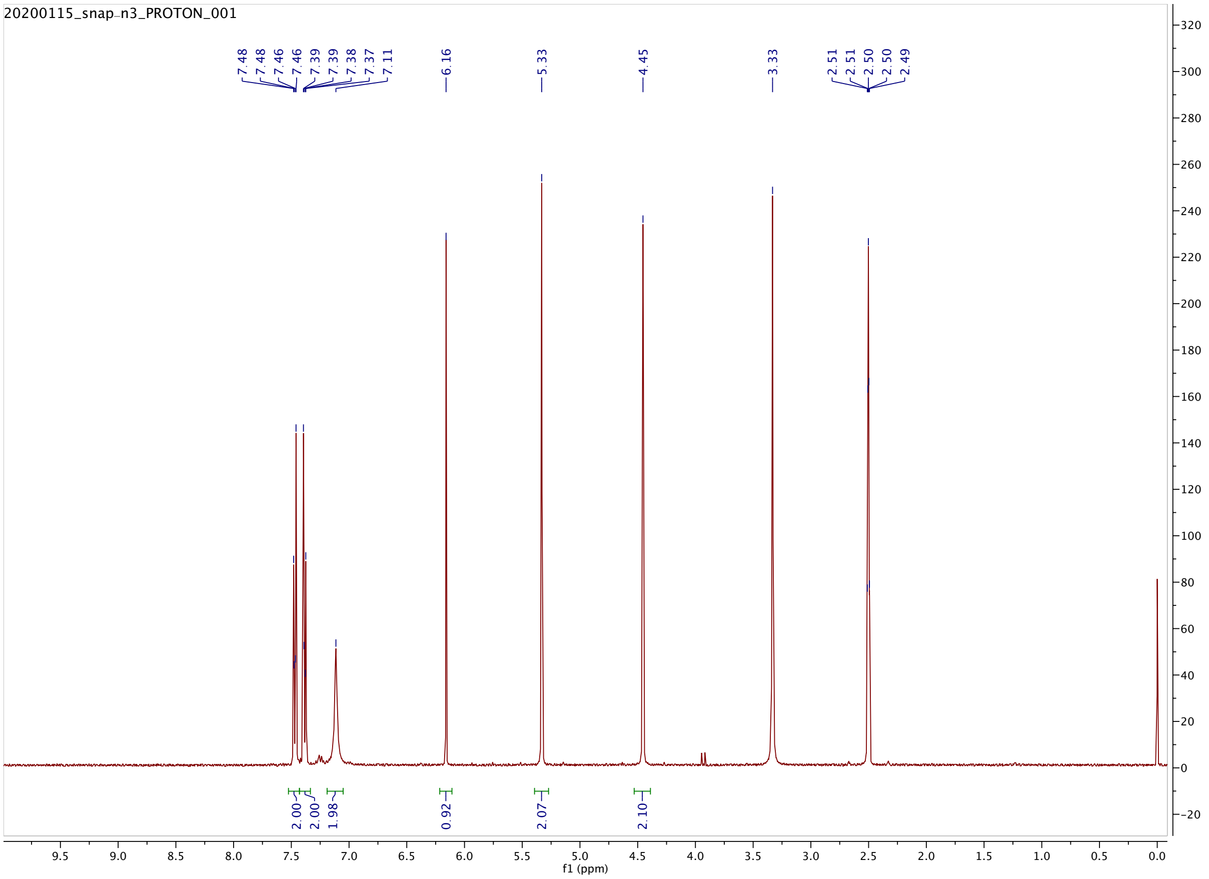
20200115_snap-n3_PROTON_001
7.48 7.48 7.46 7.46 7.39 7.39 7.38 7.37 7.11
6.16
5.33
4.45
3.33
2.51 2.51 2.50 2.50 2.49
2.00
2.00
1.98
0.92
2.07
2.10
f1 (ppm)

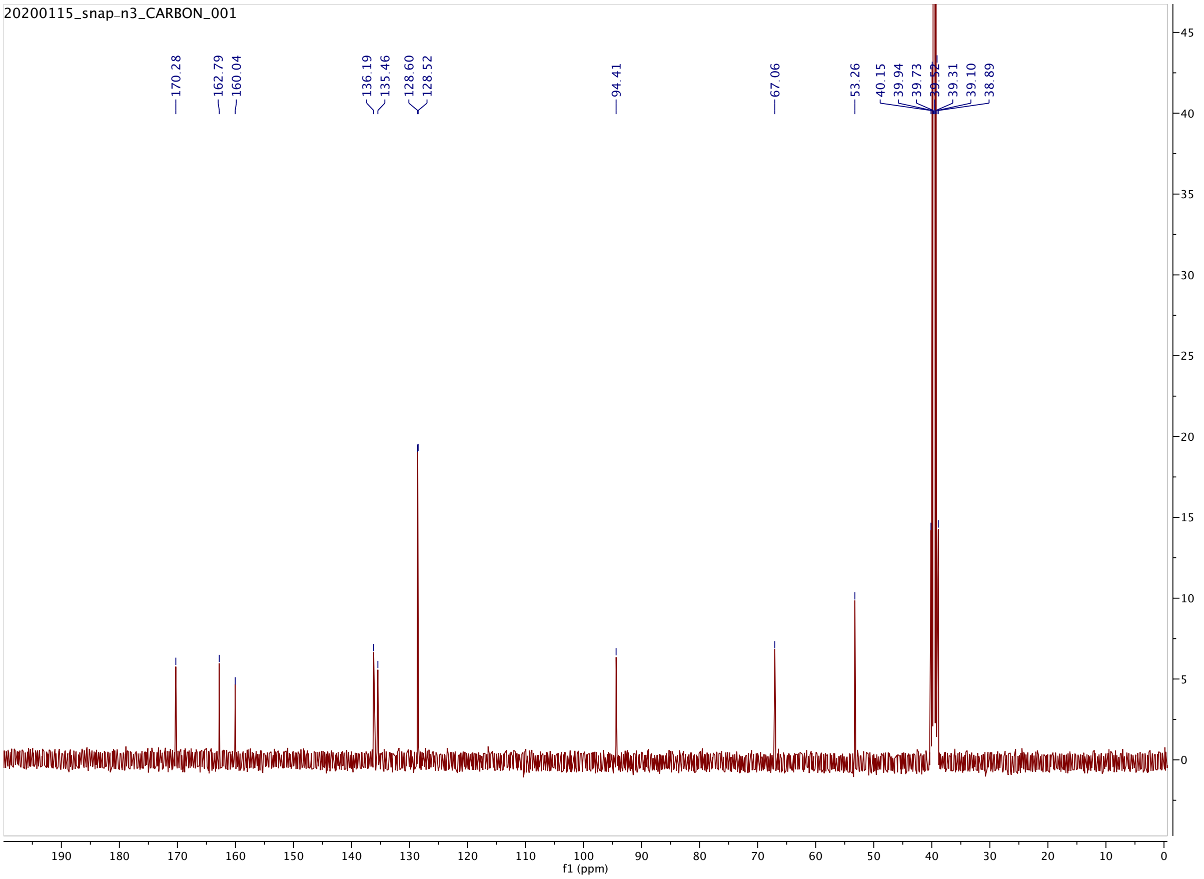
20200115_snap-n3_CARBON_001
170.28
162.79
160.04
136.19
135.46
128.60
128.52
94.41
67.06
53.26
40.15
39.94
39.73
39.52
39.31
39.10
38.89
f1 (ppm)

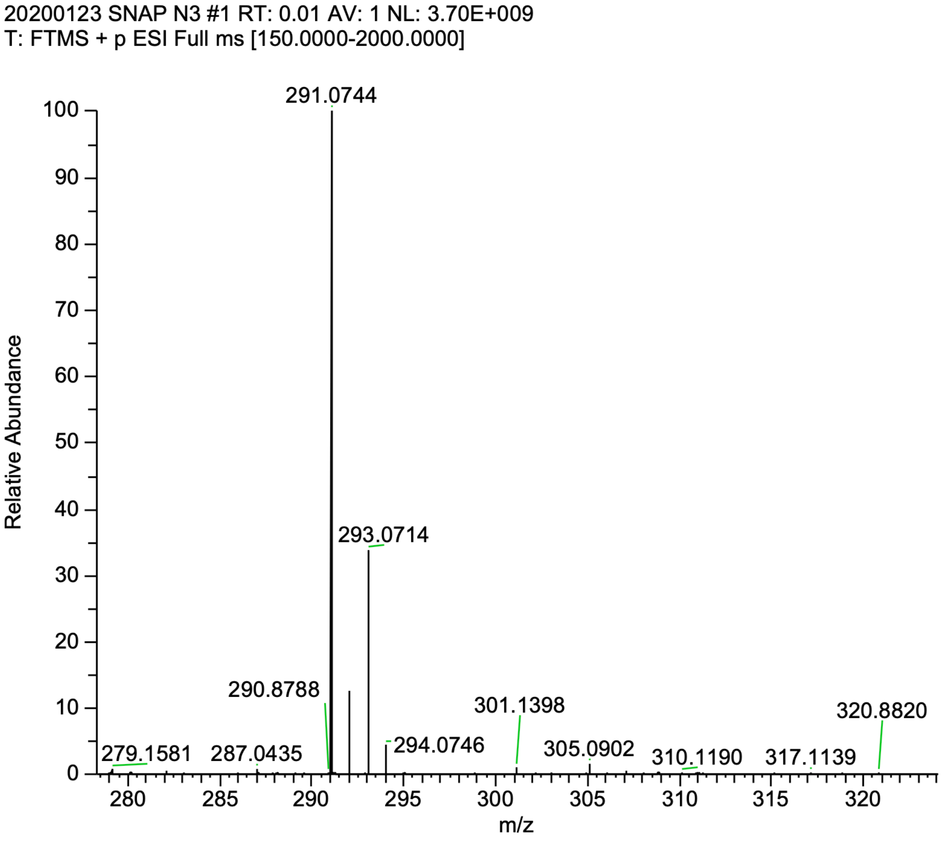


**4-((4-(aminomethyl)benzyl)oxy)-6-chloropyrimidin-2-amine (SNAP-$NH_2$). 2** (500 mg, 1.72 mmol) was dissolved into THF (60 mL) with triphenylphospine (903 mg, 3.45 mmol). Three mL of $H_2O$ were added, and the reaction was allowed to stir overnight at room temperature. The following day, all solvent was removed using a rotary evaporator and the residue was further dried under vacuum. The solid was suspended into cold DCM (~15 mL) and filtered to remove triphenyl phosphine oxide, yielding a fluffy white solid (243 mg, 57% yield).

$^1$H NMR (400 MHz, DMSO-$d_6$) δ 7.34 (q, $J$ = 8.3 Hz, 4H), 7.10 (s, 2H), 6.13 (s, 1H), 5.28 (s, 2H), 3.70 (s, 2H), 1.82 (s, 2H). $^{13}$C NMR (100 MHz, DMSO-$d_6$) δ 170.34, 162.79, 159.99, 144.32, 133.96, 128.26, 127.04, 94.42, 67.38, 45.40. HRMS (M + $H^+$): Expected: 265.0851, Found: 265.0842.

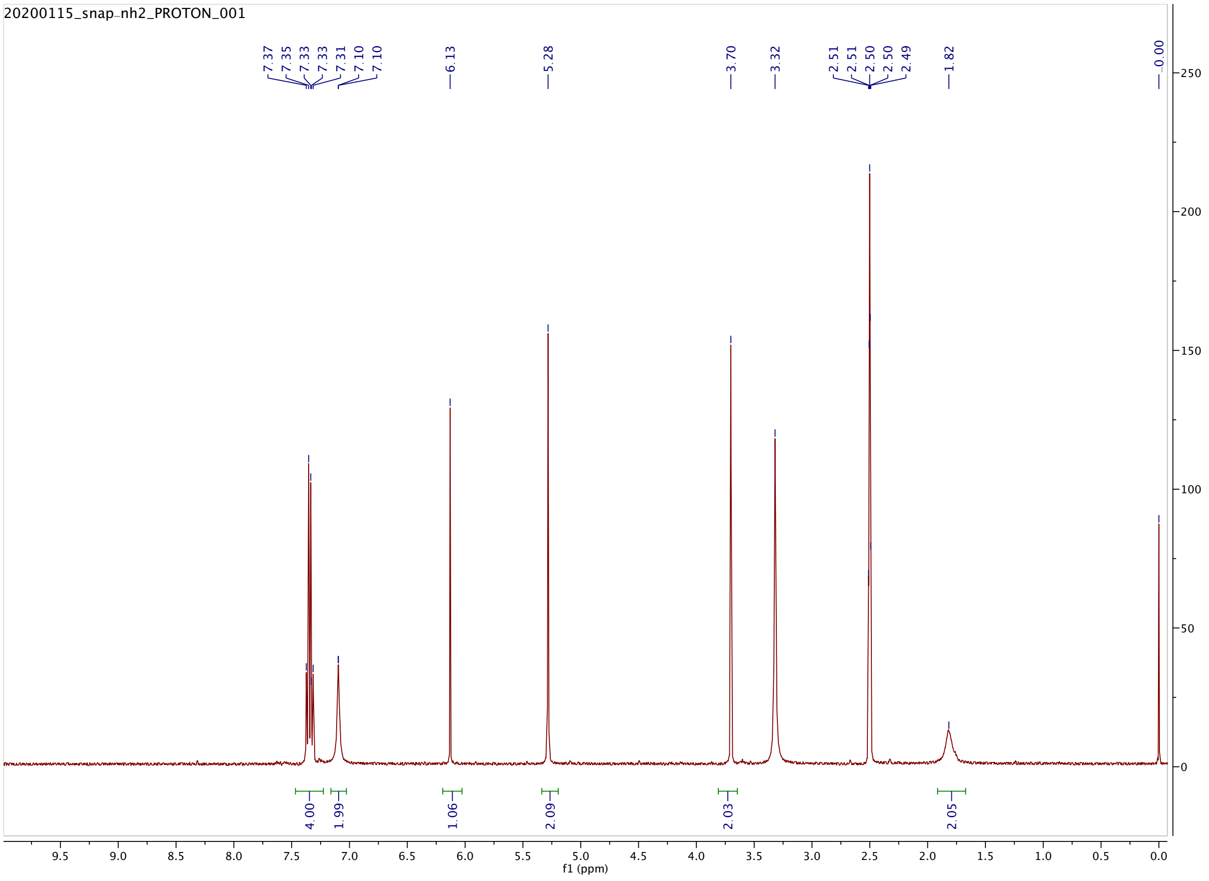
20200115_snap-nh2_PROTON_001
7.37
7.35
7.33
7.33
7.31
7.10
7.10
6.13
5.28
3.70
3.32
2.51
2.51
2.50
2.50
2.49
1.82
0.00
4.00
1.99
1.06
2.09
2.03
2.05
f1 (ppm)

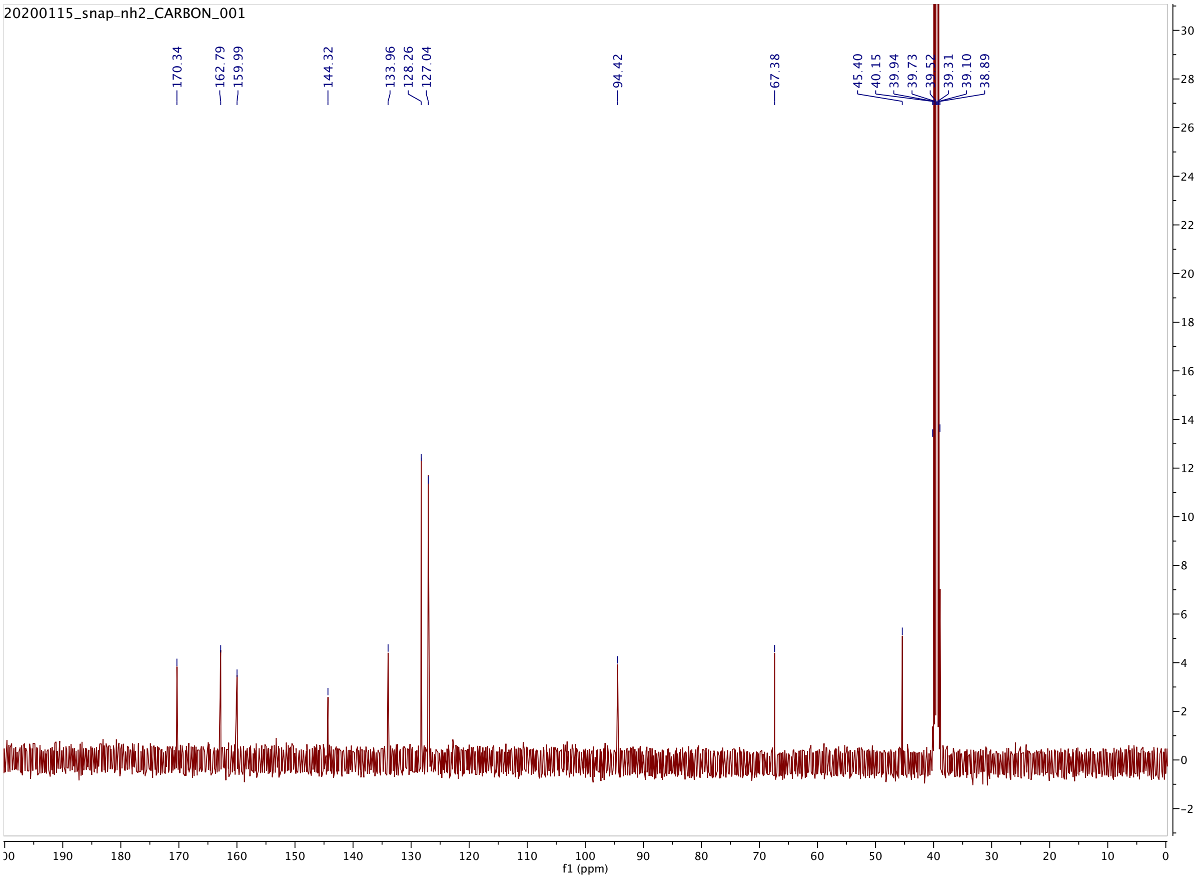


20200123 SNAP NH2 #2 RT: 0.02 AV: 1 NL: 3.30E+009
T: FTMS + p ESI Full ms [150.0000-2000.0000]

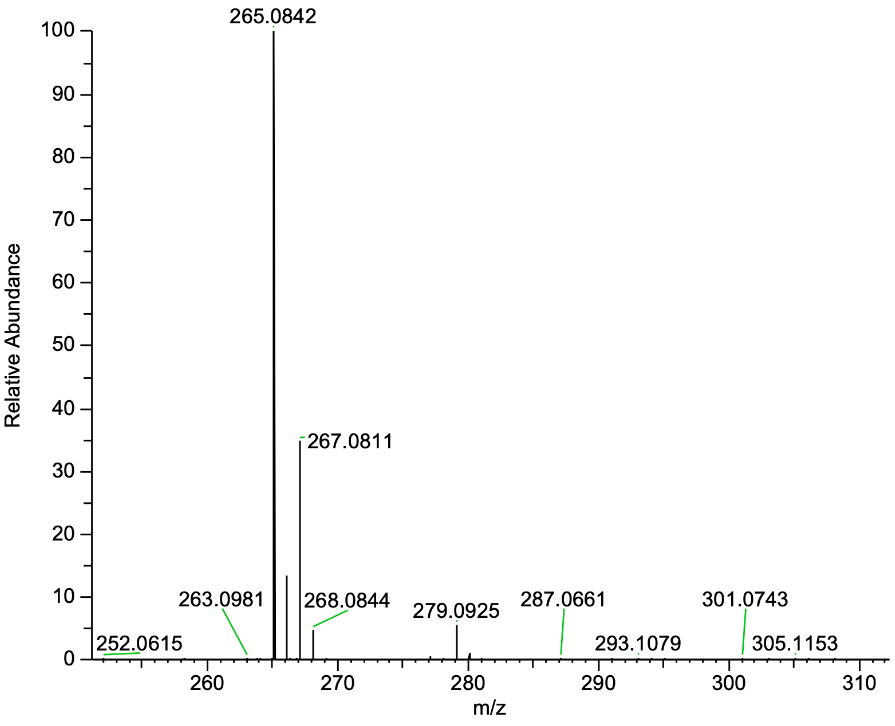

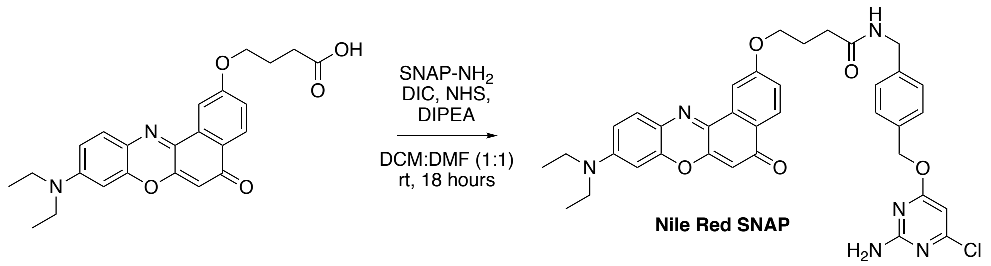


**Nile Red SNAP** was prepared by coupling SNAP-$NH_2$ to Nile Red derivatized with a carboxylic acid handle at the 9-position. Carboxy-Nile Red was prepared using a previously reported synthesis.[1] 3.5 mg (8.3 µmol) of Nile Red-CO2H was coupled to SNAP-$NH_2$ (2.6 mg, 10 µmol) using DIC (1.6 µL, 10 µmol), NHS (1.4 mg, 12.5 µmol), and DIPEA (2.9 µL, 16.7 µmol), and was allowed to stir in a 1:1 mixture of DCM (1 mL) and DMF (1 mL) overnight. The next day, the reaction was diluted into DCM, and washed with 10% citric acid, saturated $NaHCO_3$ and brine. Then the organic layer was dried with $MgSO_4$. The crude product was dried onto silica gel and purified using a gradient of 0-5% MeOH in DCM. A second column using a gradient of 10-100% EtOAc in Hexanes was necessary to completely purify the dye for subsequent photophysical characterization and cellular studies. After purification, 3.2 mg of NR-SNAP was obtained (58% yield).

$^1$H-NMR ($CDCl_3$ + Trace MeOD for solubility, 400 MHz): δ 8.15 (d, *J* = 8.7 Hz, 1H), 7.99 (d, *J* = 2.6 Hz, 1H), 7.58 (d, *J* = 9.1 Hz, 1H), 7.26 – 7.16 (m, 4H), 7.06 (dd, *J* = 8.7, 2.6 Hz, 1H), 6.65 (dd, *J* = 9.1, 2.7 Hz, 1H), 6.44 (d, *J* = 2.7 Hz, 1H), 6.28 (s, 1H), 6.07 (s, 1H), 5.18 (s, 2H), 4.41 (d, *J* = 5.2 Hz, 2H), 4.19 (t, *J* = 5.8 Hz, 2H), 3.46 (q, *J* = 7.1 Hz, 4H), 2.48 (t, *J* = 7.1 Hz, 2H), 2.22 (p, *J* = 6.2 Hz, 2H), 1.25 (t, *J* = 7.0 Hz, 6H). $^{13}$C-NMR ($CDCl_3$ + Trace MeOD for solubility, 100 MHz): δ 183.59, 172.58, 172.50, 170.88, 161.63, 160.85, 152.34, 151.04, 147.01, 139.64, 138.38, 135.32, 134.20, 131.21, 128.49, 128.03, 127.86, 125.71, 124.88, 118.36, 109.89, 106.66, 105.17, 97.23, 96.36, 67.96, 67.26, 45.22, 43.32, 32.90, 25.22, 12.71. HRMS (M + $H^+$): Expected: 667.2430, Found: 667.2429

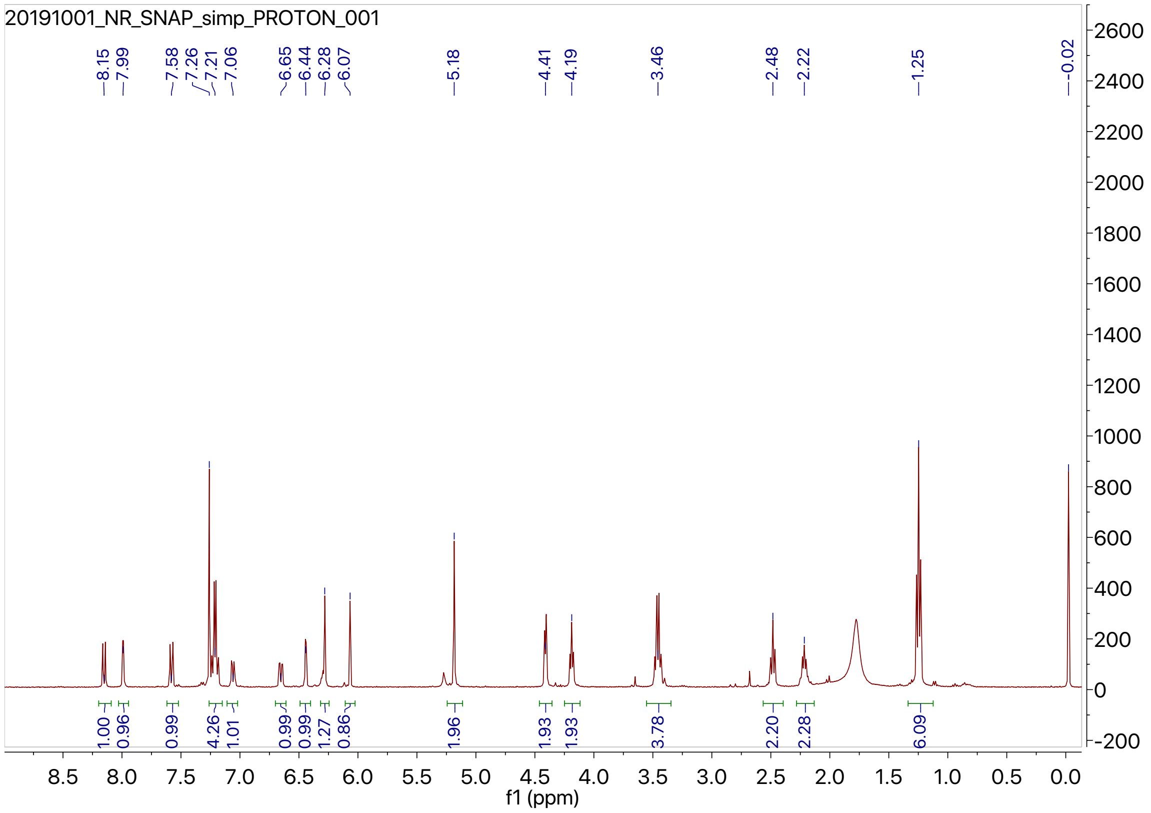
20191001_NR_SNAP_simp_PROTON_001
f1 (ppm)

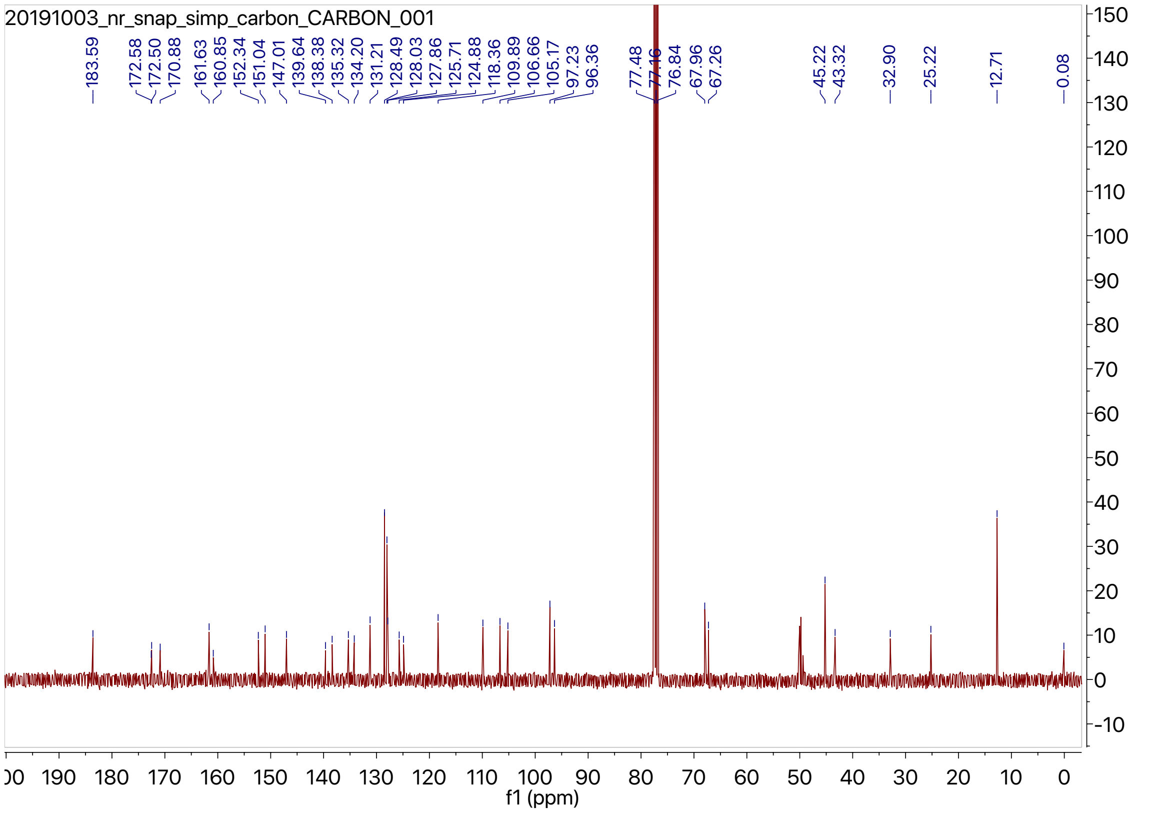
20191003_nr_snap_simp_carbon_CARBON_001
f1 (ppm)

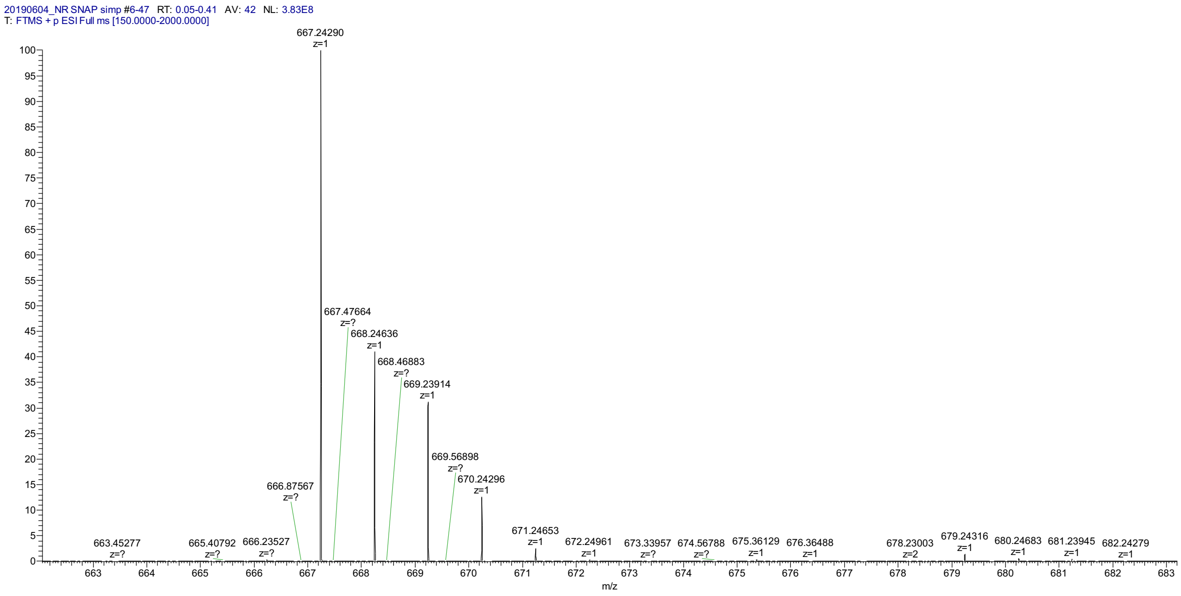
20190604_NR SNAP simp #6-47 RT: 0.05-0.41 AV: 42 NL: 3.83E8
T: FTMS + p ESI Full ms [150.0000-2000.0000]
667.24290 z=1
667.47664 z=?
668.24636 z=1
668.46883 z=?
669.23914 z=1
669.56898 z=?
670.24296 z=1
666.87567 z=?
663.45277 z=?
665.40792 z=?
666.23527 z=?
671.24653 z=1
672.24961 z=1
673.33957 z=?
674.56788 z=?
675.36129 z=1
676.36488 z=1
678.23003 z=2
679.24316 z=1
680.24683 z=1
681.23945 z=1
682.24279 z=1
m/z
